# A phosphorylation switch in PAGE4 drives MED12-mutant fibroid pathogenesis

**DOI:** 10.64898/2026.03.31.715552

**Authors:** Yih Tyng Bong, Xiaonan Liu, Yaxin Jing, Iftekhar Chowdhury, Eevi Kaasinen, Zenglai Tan, Dicle Malaymar Pinar, Zixian Wang, Auli Karhu, Niko Välimäki, Tiina Öhman, Gong-Hong Wei, Salla Keskitalo, Lauri A. Aaltonen, Markku Varjosalo

## Abstract

Recurrent somatic mutations in MED12, found in ∼70% of uterine leiomyomas (ULs), define the dominant molecular subtype of these highly prevalent tumors, yet the downstream effector mechanisms remain poorly understood.

Using an integrated multi-omics workflow, encompassing discovery-phase DDA proteomics of matched leiomyoma–myometrium pairs, validation-phase DIA proteomics across genetically stratified cohorts (MED12 p.G44D, RAD51B–HMGA2, IRS4/FH subtypes), phosphoproteomics, immunohistochemistry, and AP-MS interactomics, we identify prostate-associated gene 4 (PAGE4) as a central effector of MED12-mutant UL pathogenesis.

PAGE4 emerged as one of the most significantly upregulated proteins in MED12-mutant ULs and harbored the highest-occupancy hyperphosphorylation sites (T51, T85) in the tumor phosphoproteome, a pattern confirmed by immunohistochemistry and orthogonal phosphoproteomics. Kinase-substrate enrichment nominated HIPK2 as the primary upstream kinase, with supporting evidence from the broader CMGC family. Subsequent analysis revealed that phosphorylation acts as a molecular switch, fundamentally restructuring the PAGE4 interactome to favor the Mediator complex and RNA Pol II transcriptional machinery. This rewiring was functionally validated by ChIP-seq and luciferase reporter assays, which demonstrated corresponding shifts in transcription factor occupancy and downstream pathway activity.

Collectively, these data establish phospho-PAGE4 as a critical mechanistic node downstream of MED12 mutation, expand PAGE4 biology from prostate cancer to female reproductive tumors, and nominate it as a mechanistic biomarker and candidate therapeutic vulnerability in the most prevalent molecular subtype of uterine leiomyomas.

## Introduction

Uterine leiomyomas (ULs), commonly known as fibroids, are the most prevalent type of non-malignant pelvic tumors, with an incidence exceeding 70% among women of reproductive age. (Al-Hendy *et al*, 2017; Baird *et al*, 2003; Buyukcelebi *et al*, 2023a; Wise & Laughlin-Tommaso, 2016) These tumors originate from the smooth muscle cells of the uterine myometrium and can lead to a range of debilitating symptoms such as abnormal uterine bleeding, severe pelvic pain, reproductive complications including infertility, and pressure symptoms affecting the bladder and bowel (Tu *et al*, 2023). These symptoms significantly impair quality of life and are commonly treated with surgical interventions, including hysterectomy and myomectomy, both of which compromise future fertility and carry significant risks (Bano *et al*, 2023; Yu *et al*, 2022). In the United States alone, ULs account for approximately 200,000 hysterectomies annually and impose an economic burden exceeding $34 billion per year (Cardozo *et al*, 2012). Although recent advances in pharmacological approaches for management of ULs, such as selective progesterone receptor modulators, GnRH antagonists and more recently linzagolix (Ali *et al*, 2023; Donnez *et al*, 2022), have shown promising outcomes in reducing fibroid size and alleviating symptoms, these therapies remain largely temporary and are often used as adjuncts or bridges to surgical intervention. Consequently, the incomplete understanding of UL pathophysiology remains a critical barrier to developing definitive, non-invasive therapies (El Sabeh *et al*, 2021).

Extensive research has established that approximately 70% of ULs harbor somatic mutations in the Mediator Complex Subunit 12 (*MED12*) gene (Mäkinen *et al*, 2011). These alterations are highly recurrent and almost exclusively cluster within exon 2, most prominently at codon 44, where substitutions such as the recurrent p.G44D arise(Mäkinen *et al*., 2011). MED12 is a critical component of the Mediator complex, a multiprotein assembly that bridges transcription factors with RNA polymerase II to regulate gene expression (Philibert & Madan, 2007). In addition to initiating transcription, the Mediator complex coordinates chromatin remodeling and integrates diverse signaling inputs into coherent transcriptional programs (Allen & Taatjes, 2015). In our previous work we identified mutations of *MED12* with widespread transcriptional dysregulation and impaired Mediator associated kinase activity (Turunen *et al*, 2014a). Despite this genetic insight, the molecular mechanisms linking MED12 mutations to fibroid pathogenesis remain incompletely understood. Most studies have focused on transcriptomic analyses, which, while informative, provide only a partial view given the well-documented disconnect between mRNA and protein abundance (Greenbaum *et al*, 2003; Liu *et al*, 2016). Furthermore, while pioneering proteomic studies have identified candidate proteins (Ciebiera *et al*, 2024; Jamaluddin *et al*, 2018a), these efforts have been limited in scope and have not systematically integrated proteomics with functional validation. A comprehensive proteomic approach that captures both protein expression changes and post-translational modifications, particularly phosphorylation, a key regulatory mechanism, is essential to bridge the gap between *MED12* genotype and UL phenotype.

To address this knowledge gap and establish the molecular consequences of *MED12* mutations, we employed a multi-stage proteomic and functional genomics strategy. We began with unbiased data-dependent acquisition (DDA) mass spectrometry, ideal for comprehensive profiling of the proteome and phosphoproteome in matched leiomyoma-myometrium tissue pairs. Key findings were validated using immunohistochemistry. To broaden the analysis and ensure quantitative robustness across the major molecular subtypes of ULs, we next applied data-independent acquisition (DIA) mass spectrometry to an expanded cohort representing MED12-mutant tumors, high mobility group AT-hook 2 (HMGA2)-activated tumors driven by *RAD51B* enhancer rearrangements (approximately 10-15%), and rare fumarate hydratase (FH)-deficient subgroup (approximately 0.4-1.6%)(Mäkinen *et al*., 2011). This DIA cohort also included tumors with collagen, type IV, alpha 5 and collagen, type IV, alpha 6 (COL4A5-COL4A6) deletions (approximately 2%), a genomic alteration that has been associated with a distinct transcriptional profile characterized by strong IRS4 upregulation and has been proposed to define a rare fourth UL subgroup (Mehine *et al*, 2016; Mehine *et al*, 2013). In our cohort, IRS4 overexpression was observed in tumors carrying COL4A5-COL4A6 deletions on a concurrent MED12-mutant background. Accordingly, these samples were analyzed together as a unified COL4A5-COL4A6 subgroup for downstream proteomic analyses. DIA proteomics enabled reproducible, high-resolution quantification of both protein abundance and phosphosite occupancy normalized to parent protein levels (Di *et al*, 2023).

This integrative analysis convergently identified prostate-associated gene 4 (PAGE4) as the most compelling candidate linking MED12 mutation to uterine fibroid pathogenesis. PAGE4 was among the most strongly upregulated proteins in MED12-mutant tumors, displayed extensive tumor-specific hyperphosphorylation, and showed a consistent association with the MED12-mutant subtype across all analytical platforms. While PAGE4 is recognized as a biomarker for male-specific diseases, particularly prostate cancer (Lin *et al*, 2018b) where its overexpression is associated with aggressive tumor behavior (Kulkarni *et al*, 2016), its function in female reproductive biology has remained unexplored. Through systematic interactome mapping using complementary affinity purification and proximity labeling approaches, we discovered that PAGE4 phosphorylation acts as a molecular switch, dramatically remodeling its protein interaction landscape to favor associations with the Mediator complex and core transcriptional machinery. These phosphorylation-dependent interactions were functionally validated through ChIP-seq analysis of RNA polymerase II occupancy and luciferase-based transcriptional reporter assays, which revealed corresponding changes in transcription factor binding patterns and pathway activation. Our findings establish phospho-PAGE4 as a central mechanistic node in MED12-mutant fibroid pathogenesis, expand the biological role of PAGE4 beyond prostate cancer to encompass female reproductive tract neoplasia, and identify a promising biomarker and potential therapeutic target for the most common molecular subtype of ULs.

## Results

### Proteomic profiling reveals a distinct molecular phenotype in MED12 G44D Leiomyomas

To investigate the molecular basis of UL harboring the MED12 G44D mutation, we performed a comprehensive proteomic analysis using paired UL and adjacent myometrium (MM) tissue samples (**Figure 1A**). These were collected with written informed consent from nine patients during hysterectomy or myomectomy, and all data were generated using data-dependent acquisition (DDA) proteomics. Principal component analysis (PCA) using the expression levels of all proteins separated the two types of samples relatively well (**Figure 1B**), with the first component explaining nearly 33.9% of the variation, while the second component explaining 12.3%, highlighting that UL results in substantial proteomic differences. Comparative analysis revealed 161 proteins significantly upregulated and 70 downregulated in UL versus MM (p-value < 0.05, log2 FC > 2) (**Figure 1C; Table S1**). Leiomyomas were characterized by a pronounced upregulation of extracellular matrix (ECM) and adhesion components, including COL1A1, COL3A1, POSTN, and the matrix-crosslinking enzyme PLOD1 (**Figure 1C**). This fibrotic signature was accompanied by increases in RNA-processing factors (EIF4A3, PRPF19, SF3B6) and proteins involved in secretion and stress-response (HYOU1), suggesting a state of heightened RNA metabolism and secretory activity (Jamaluddin *et al*, 2018b; Rafique *et al*, 2017). Conversely, proteins critical for smooth muscle contractility (TPM2), cytoskeletal integrity (SPTAN1, SPTBN1), and cell-cell adhesion (CDH13, TJP1) were significantly downregulated (**Figure 1C**). This coordinated shift indicates a fundamental phenotypic transition, where UL tumors gain pro-fibrotic and RNA-handling capabilities while losing the contractile identity of the native myometrium (Ura *et al*, 2019).

**Figure 1:**
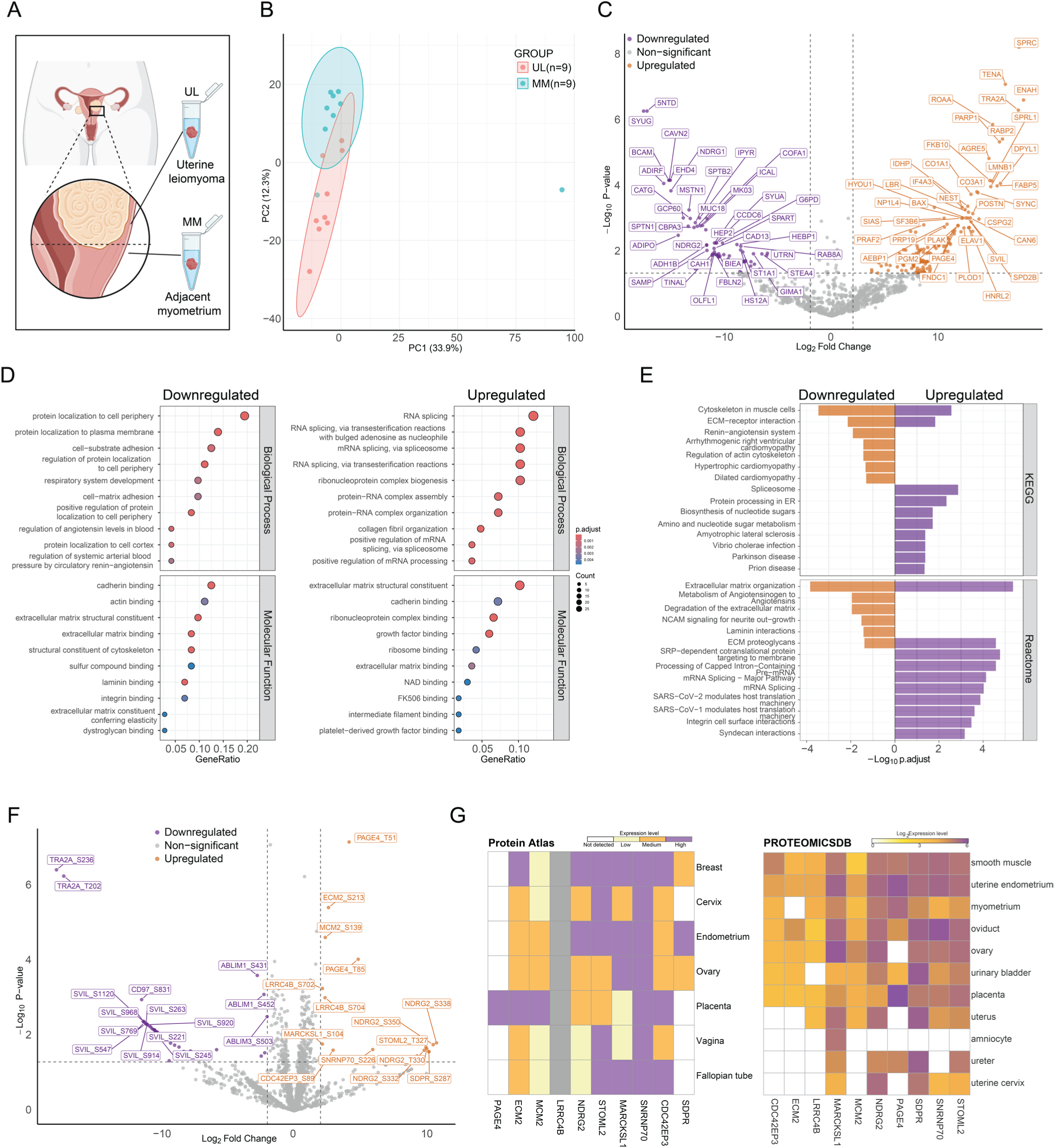
Proteomic and phosphoproteomic profiling identifies PAGE4 as a potential marker in ULs. (**A**) Schematic of the study depicted the collection of matched uterine leiomyoma (UL) and adjacent myometrium (MM) tissues during surgery, followed by their processing for omics analysis. (**B**) Principal component analysis (PCA) of the global proteome distinguishes UL from MM tissues. PC1 accounts for 33.9% and PC2 for 12.3% of the total variance. Individual samples are shown as points with 95% confidence ellipses for each group (UL n=9; MM n=9 matched patient pairs). (**C**) Volcano plot of differential protein abundance between UL and MM. Significantly changing proteins (|log₂ fold-change| > 2, p < 0.05, two-sided t-test) are highlighted in orange (upregulated in UL) or purple (downregulated in UL). Selected protein names are labeled. (**D**) GO enrichment analysis of differentially abundant proteins. Dot size indicates the number of proteins in each category; color represents adjusted p-value. (**E**) Pathway enrichment analysis of up- and downregulated proteins using Reactome and KEGG databases. Bar height corresponds to statistical significance (-log₁₀ adjusted p-value). Upregulated proteins associate with ECM organization and muscle contraction; downregulated proteins associate with spliceosome and translation machinery. (**F**) Volcano plot of differentially occupied phosphorylation sites (UL vs. MM). Significantly changing phosphosites (|log₂ fold-change| > 2, p < 0.05) are highlighted. PAGE4 phosphosites T51 and T85 are among the most significantly hyperphosphorylated in UL (indicated). (**G**) Cross-validation of protein expression patterns using external databases. Heatmaps show expression levels from the Human Protein Atlas (categorical) and ProteomicsDB (log-transformed expression values) for selected proteins identified in panels (C) and (F).

Gene Ontology (GO) enrichment analysis substantiated these proteomic trends, delineating a clear functional partition (**Figure 1D**). Biological processes downregulated in UL were dominated by terms related to cell cortex organization, cell-matrix adhesion, and the regulation of vascular tone, underscoring a loss of smooth muscle homeostasis. Molecular functions were depleted for cadherin, intermediate filament, and ECM binding. Conversely, upregulated processes in UL were overwhelmingly associated with RNA splicing, ribonucleoprotein complex assembly, and collagen fibril organization. The corresponding molecular functions highlighted integrin and laminin binding, driven by the accumulated ECM and adhesion molecules (Jamaluddin *et al*, 2019; Jamaluddin *et al*., 2018b), pointing to active matrix deposition and cell-ECM engagement.

Pathway analysis using KEGG and Reactome provided further mechanistic context for this phenotypic switch (**Figure 1E**). The pathways downregulated in UL consistently highlighted a suppression of smooth muscle function, including “cytoskeleton in muscle cells,” “regulation of actin cytoskeleton,” and various cardiomyopathy terms, alongside a notable reduction in “extracellular matrix organization.” This suggests a dismantling of the native, organized contractile apparatus. In contrast, upregulated pathways revealed a tumor focused on post-transcriptional control and metabolism, with strong enrichment for the “spliceosome” and “SRP-dependent co-translational protein targeting to membrane.” This was accompanied by specific gains in “integrin cell surface interactions” and “syndecan interactions,” underscoring an active reprogramming of the cell-surface interface.

Together, these findings underscore a proteome-wide shift in gaining RNA processing, translational capacity, and specific cell-ECM communication while losing the contractile and structural programs characteristic of normal myometrium. Crucially, the involvement of dynamic, kinase-driven pathways, such as the “regulation of actin cytoskeleton” and “integrin cell surface interactions”, suggested that protein abundance alone could not fully explain the mechanistic drivers of this phenotypic shift. This gap in understanding directly motivated our subsequent phosphoproteomics analysis to identify the critical phosphorylation events coordinating these cytoskeletal and RNA-processing changes.

### Phosphoproteomic analysis identifies aberrant kinase networks and PAGE4 hyperphosphorylation

To delineate the post-translational drivers of the phenotypic shifts observed in MED12 G44D-mutant leiomyomas, we advanced our investigation by analyzing the phosphoproteome using paired tissue samples (**Figure 1F; Table S1**). Among the most significantly hyperphosphorylated sites in ULs were PAGE4 at Thr51 and Thr85, alongside sites on proteins regulating transcription and RNA splicing (MCM2_S139, SNRNP70_S226), cytoskeletal and adhesion dynamics (MARCKSL1_S104, ECM2_S213), and signaling modules (NDRG2 cluster, LRRC4B_S702/S704). Conversely, a broad pattern of hypophosphorylation was observed on proteins critical for cytoskeletal integrity and contractility, including multiple sites on the actin-crosslinker SVIL and the adhesion complex component ABLIM1. Interestingly, PAGE4, traditionally recognized as a male-specific marker in prostate cancer and other urogenital diseases, stood out as a prominently hyperphosphorylated protein in these female-specific tumors, suggesting a previously unrecognized, sex-independent role in cellular signaling and tumor progression.

We next sought to prioritize candidates for mechanistic follow-up. Because phosphorylation reflects kinase signaling activity and downstream programs that govern transcription, cytoskeletal remodeling, and extracellular matrix deposition, which are processes central to tumorigenesis and fibrosis, we prioritized the most strongly altered phosphosites as likely proximal regulators of the UL phenotype. To ensure robustness and translational relevance, we focused on proteins that were concurrently hyperphosphorylated and upregulated in UL (**Figure 1C and 1F**). This simple two-step screen favors candidates with coherent regulation, improves their detectability for follow-up assays and biomarkers, and fits the fibrotic, secretory phenotype we observe in UL. To place these hits in tissue context, we checked baseline expression in public proteomic resources (**Figure 1G**). A tissue-level qualitative view from the Protein Atlas and a quantitative profile from ProteomicsDB confirmed that several proteins carrying UL-hyperphosphorylated sites, are endogenously expressed in female reproductive tissues. This orthogonal data supported the biological relevance of our phosphosite findings and reinforced the prioritization of targets for downstream validation.

However, not all high-signal candidates were pursued equally. We de-prioritized ubiquitous cell-cycle and splicing factors (e.g., MCM2, SNRNP70), whose phosphorylation typically reflects proliferation rather than a UL-specific mechanism. Broad cytoskeletal adaptors (e.g., SVIL, ABLIM1) and secreted ECM-related proteins (e.g., ECM2, SDPR) likely reflect the global shift toward fibrosis already evident at the pathway level, making it harder to tie them to MED12 p.G44D in a focused way. We also set aside stress-response nodes such as NDRG2, which participate in pleiotropic signaling and would complicate a clean mechanistic story in this first pass.

By contrast, PAGE4 offered a rare combination of features: it is upregulated in UL at protein level, harbors two of the most significant UL-hyperphosphorylated sites (T51/T85), is detectably expressed in uterine tissues. This was a surprising finding, as PAGE4 has been defined almost exclusively as a male-specific cancer/testis antigen, a characterization supported by its high expression in prostate and testis and its near absence in normal female tissues (**Figure S1**). Its clear involvement in uterine leiomyomas uncovers a novel, sex-independent role for PAGE4, suggesting its pathological reactivation in female reproductive tissues through stress or epigenetic remodeling.

### Immunohistochemistry establishes PAGE4 staining elevation in UL

Building on our proteomic identification of PAGE4 as a potential candidate protein marker for ULs, we next assessed its expression pattern in patient tissues by immunohistochemistry (IHC) (**Figure 2A**). Because the proteomic discovery was driven by *MED12*-mutant tumors, the IHC analysis focused specifically on *MED12*-mutant ULs and their matched myometrial controls. In addition to PAGE4, we also stained for progesterone receptor (PR), and estrogen receptor (ER), being routinely used diagnostic markers for ULs in clinical pathology (Leitao *et al*, 2004). PAGE4 staining was consistently absent or very weak in myometrium but strong and diffuse in MED12 UL tissues, localizing predominantly to the cytoplasm. In contrast, ER and PR staining patterns were comparable between MM and UL, showing variable nuclear positivity without consistent upregulation (**Figure 2A**), highlighting previously noted complex relationship between their levels and UL (Mehasseb *et al*, 2011). Quantitative image analysis confirmed that PAGE4 displayed significantly higher staining intensity in ULs compared to MM (p < 0.05), while ER and PR did not differ significantly between tissue types (**Figure 2B**), further complicating their reliability as consistent immunohistochemical markers (Jakimiuk *et al*, 2004). To further validate these findings at the transcript level, we re-analyzed published uterine fibroid RNA-seq data (Berta *et al*, 2021). The result highlights *PAGE4* among the most strongly upregulated genes in MED12 UL compared with paired MM, providing independent transcriptomic support for our proteomic and IHC observations (**Figure 2C**).

**Figure 2:**
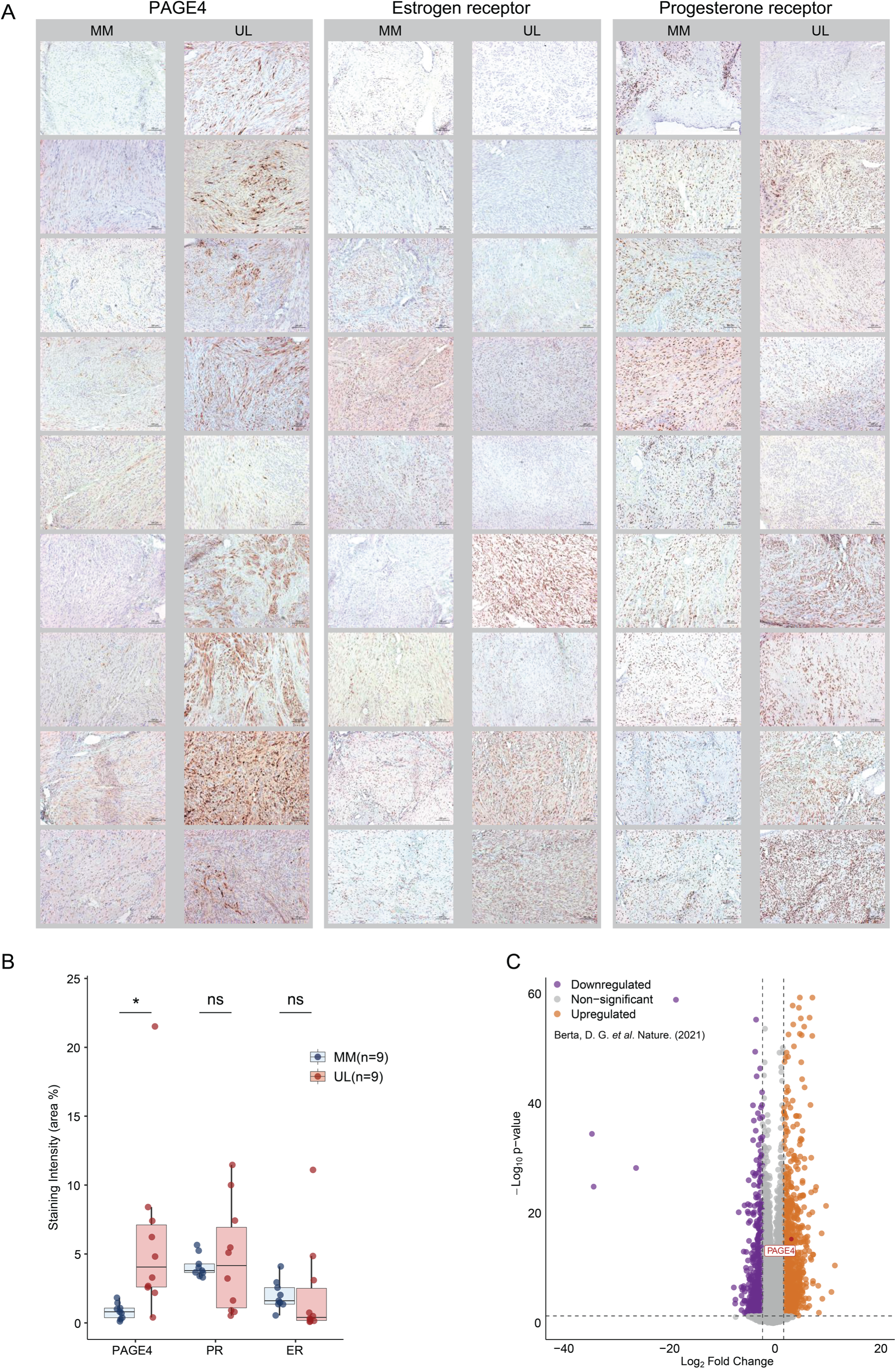
Immunohistochemical validation of PAGE4, PR, and ER expression in *MED1*2 ULs. **(A)** Representative immunohistochemical staining of PAGE4, estrogen receptor (ER), and progesterone receptor (PR) in nine matched pairs of myometrium (MM, left columns) and MED12 ULs (right columns) tissues. Each row represents tissue from an individual patient, illustrating inter-patient variability in hormone receptor and PAGE4 expression. Scale bar: 100 µm. (**B)** Quantification of staining intensity expressed as percentage of positive area. Box plots display median (center line), interquartile range (box), and individual data points for MM (n=9) and UL (n=9) matched tissues. PAGE4 expression is significantly elevated in UL compared to MM (*p ≤ 0.05, paired t-test), while PR and ER show no significant differences (ns, p > 0.05). (**C)** Transcriptomic validation using published RNA-seq data (Berta et al., Nature 2021). Volcano plot shows differential gene expression between MED12 UL (n=18) and MM (n=9) samples. PAGE4 is among the significantly upregulated genes in UL tissues (gray box), corroborating our proteomic findings in an independent patient cohort.

Together, these data establish that PAGE4 is a robustly induced protein in MED12 ULs, distinguishable from surrounding normal tissue, and suggest that PAGE4 may complement or outperform conventional hormone-receptor markers (ER, PR) in capturing the molecular identity of MED12 ULs.

### System-level protein and phosphorylation patterns differentiate UL subclasses

To validate the initial PAGE4-associated signatures identified in our MED12 p.G44D discovery cohort using DDA proteomics, and to determine whether these signatures were reproducible across genetically distinct uterine leiomyoma subclasses rather than being cohort-dependent, we expanded the analysis using data-independent acquisition (DIA) proteomics. While the discovery phase focused on matched MED12-mutant leiomyoma-myometrium pairs, the DIA cohort was designed to capture the major molecular contexts of ULs and included four groups: MED12-mutant tumors (n=9 pairs), HMGA2-activated tumors containing *RAD51B* enhancer rearrangements (n =10 pairs), FH-deficient tumors (n=5 pairs), and tumors with COL4A5-COL4A6 deletions (n = 5 pairs). Previous genomic studies have shown that alterations affecting the COL4A5-COL4A6 locus are associated with a distinct transcriptional profile characterized by strong IRS4 upregulation and have been proposed to define a molecular UL subtype (Mehine *et al*., 2016; Mehine *et al*., 2013). In our cohort, all tumors classified within the COL4A5-COL4A6 group also carried concurrent MED12 mutations, indicating that this alteration can occur in combination with established UL driver events. Nevertheless, we analyzed these tumors as a separate COL4A5-COL4A6 subclass because their defining alteration and expression signature are linked to the COL4A5-COL4A6 locus and IRS4 upregulation rather than to MED12 status alone. For clarity, the four groups are hereafter referred to as the MED12, COL4A5-COL4A6, HMGA2, and FH subclasses.

Using this DIA framework, we quantified both the global proteome and phosphoproteome across all UL groups and their corresponding myometrial tissues. Unsupervised principal component analysis (PCA) consistently separated tumors from myometrium at the levels of total protein abundance, global phosphorylation, and individual phosphosite measurements. At the global proteome level (**Figure 3A**), samples formed structured clusters in which MED12-derived ULs and its corresponding MMs occupied a distinct region, separating from the other subclasses primarily along PC2. Within the MED12 cluster, UL and MM showed a consistent shift, indicating tumor-associated differences in protein abundance in this subtype. HMGA2 samples occupied an intermediate region of the PCA space, whereas COL4A5-COL4A6 and FH samples largely clustered at higher PC2 values, suggesting comparatively similar global proteome profiles between these two subclasses at the level captured by PC1-PC2. A single HMGA2_UL sample appeared as an outlier along PC1, indicating either biological heterogeneity or a sample-specific effect. Compared to the global proteome, phosphorylation-level measurements produced clearer tissue-associated structuring across multiple subclasses (**Figure 3A**). Particularly, MED12 UL samples shifted toward higher PC2 values relative to MED12 MMs, and COL4A5-COL4A6 UL samples similarly separated from their myometrial counterparts, indicating consistent tumor associated phosphoregulation. HMGA2 samples showed a pronounced separation pattern primarily along PC1, with HMGA2 ULs tending toward negative PC1 values relative to HMGA2 MMs. FH samples were more dispersed across the PCA space, with partial overlap between FH UL and FH MM, consistent with greater heterogeneity of phosphorylation states in this subclass and/or reduced separation power due to smaller cohort size. At the phosphosite level (**Figure 3A**), most MM samples clustered tightly near the origin and positive PC2 values, whereas UL samples extended away from this cluster along a trajectory toward lower PC2 values. This divergence was particularly evident for HMGA2 UL and FH UL samples, consistent with increased variability and stronger tumor-associated signaling differences at the individual phosphosite level compared to the global protein layer. Two sample-specific deviations were also apparent: one COL4A5-COL4A6 UL sample separated markedly at high PC1 and PC2 values, and one FH UL sample showed separation along PC1, highlighting inter-tumor heterogeneity that becomes more apparent when phosphorylation is resolved at individual sites.

**Figure 3:**
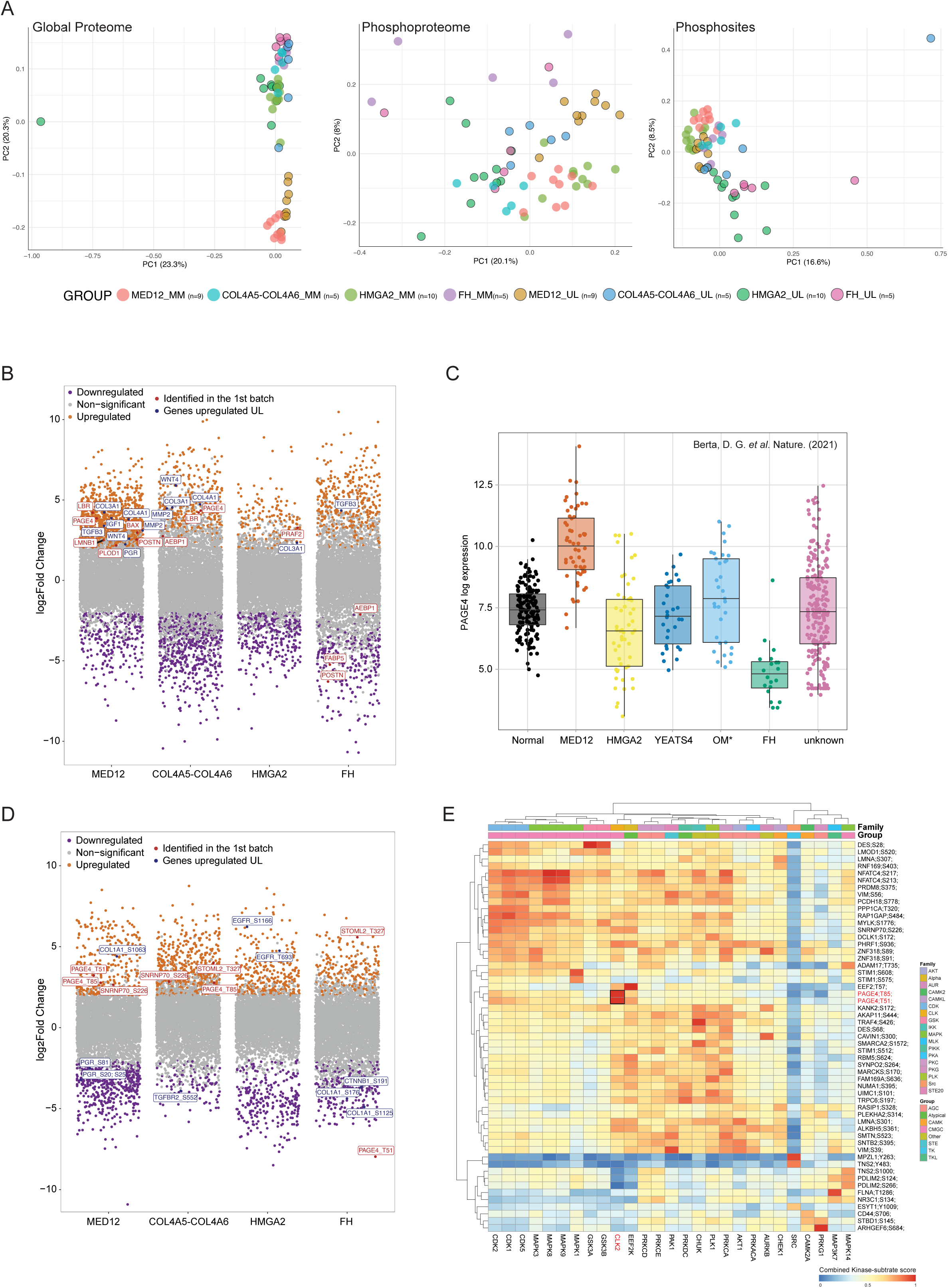
PAGE4 expression and phosphorylation are specifically enriched in MED12 ULs. (**A**) Principal component analysis (PCA) of global proteome (left), phosphoproteome (middle) and phosphosite (right) from four UL subtypes and their matched myometrium. Data were acquired using DIA mass spectrometry. Cohorts include MED12 (n=9 pairs), COL4A5-COL4A6 (n=5 pairs), HMGA2 (n=10 pairs), and FH (n=5 pairs). (**B**) Volcano plots of differentially abundant proteins (upper panels) and differentially occupied phosphorylation sites (lower panels) for each UL subtype compared to matched MM. Significantly changing features (|log₂ fold-change| ≥ 1.5, p < 0.05, two-sided t-test) are highlighted in different colors. (**C**) PAGE4 mRNA expression levels across UL molecular subtypes and normal myometrium. Data adapted from Berta et al., Nature (2021). Subtypes include: MED12, HMGA2, YEATS4, OM*, FH, and Unknown (no identified driver mutations). Box plots show median, quartiles, and individual data points. (**D**) Validation of phosphosite regulation in the expanded DIA cohort. Volcano plots show protein abundance-normalized phosphosite occupancy changes (UL vs. MM) for each genotype. Phosphosites were quantified using DIA-MS, and their abundance was normalized to changes in parent protein levels before phosphoenrichment. Significantly changing sites (|log₂ fold-change| ≥ 1.5, p < 0.05, two-sided t-test) are highlighted in different colors. (**E**) Kinase-substrate enrichment analysis identifies activated kinase families in MED12 ULs. Heatmap displays substrate enrichment scores from PhosR analysis, revealing preferential activation of CMGC family kinases. Color scale represents combined kinase-substrate enrichment score (0 to 1).

These patterns suggested that specific protein drivers contribute to both shared and subtype-dependent UL states. We therefore interrogated differential protein abundance and phosphorylation in each subclass and evaluate the specificity of PAGE4 across subclasses.

### PAGE4 expression is selectively enriched in MED12-mutant UL

At the protein level, genotype-specific volcano plots largely recapitulated key features observed in the discovery dataset while also revealing subtype-dependent differences across the expanded DIA cohort (**Figure 3B; Table S2**). COL3A1 is the only shared marker, significantly upregulated across three subtypes (MED12, COL4A5-COL4A6 and HMGA2), confirming its pan-UL role function as biomarker(Jamaluddin *et al*., 2018a).

The MED12 subclass exhibited the broadest spectrum of upregulated proteins. Notably, PAGE4 was upregulated alongside multiple markers associated with hormone signaling (WNT4, PGR, IGF1) (Bulun *et al*, 2025) and extracellular matrix remodeling (COL4A1, MMP2, TGFB3, POSTN)(Islam *et al*, 2017). Additionally, nuclear envelope-associated proteins (LMNB1, LBR) (Reilly *et al*, 2022) and PLOD1, a lysyl hydroxylase involved in collagen crosslinking, were also upregulated (**Figure 3B**). Collectively, these protein expression patterns are consistent with the amplification of fibrotic and hormone-responsive programs in this subclass(Moyo *et al*, 2020). Interestingly, PAGE4 appeared also as one of the significantly upregulated proteins in the COL4A5-COL4A6 subclasses (**Figure 3B**), whereas it was not detected in HMGA2-activated or FH-deficient tumors. Since all COL4A5-COL4A6 tumors in our cohort also carried concurrent MED12 mutations, this pattern aligns with PAGE4 elevation tracking with the MED12-mutant background. Besides PAGE4, COL4A5-COL4A6 subclass showed upregulation of several extracellular matrix markers (WNT4, COL3A1, COL4A1, and MMP2), and this subclass additionally highlighted AEBP1 and LBR which highlighted in the discovery cohort (**Figure 3B; Table S2)**. In contrast, the HMGA2 subclass displayed a more restricted set of upregulated proteins, with PRAF2 and COL3A1 emerging as the upregulated protein (**Figure 3B**). The FH subclass showed a distinct pattern in which TGFB3, a well-established leiomyoma-associated factor(Arici & Sozen, 2000), was the only prominently upregulated protein, whereas AEBP1, FABP5, and POSTN, which were increased in the MED12 discovery cohort and/or in other UL subclasses, were indicated as downregulated in FH-deficient tumors (**Figure 3B**). This divergence is notable because FH-deficient leiomyomas are generally considered a molecularly distinct driver class from canonical MED12-mutant tumors, and previous genomic studies have shown that MED12 mutations and FH inactivation are mutually exclusive in ULs (Kämpjärvi *et al*, 2016).

To further interpret protein level differences between different genotypes, we performed Gene Ontology (GO) and pathway enrichment analysis on the significantly changing proteins from each molecular subtype (p < 0.05; UL vs. MM). In the MED12 subclass (**Figure S2A**), proteins increased in UL were enriched for RNA-centric processes, consistent with what we observed in the discovery cohort (**Figure 1D**). This aligns with prior work showing that leiomyoma-associated MED12 exon 2 mutations change Mediator function, including loss of Mediator-associated CDK activity, and are linked to broad transcriptional dysregulation (Turunen *et al*, 2014b). Specifically, both MED12 and COL4A5-COL4A6 tumors showed enrichment of ECM and adhesion-related protein modules (**Figure S2A and S2B**). This aligns with our discovery cohort (**Figure 1E**) and supports previous findings that MED12-mutant fibroids exhibit distinct collagen-rich ECM and ECM-organization signatures (Ishikawa *et al*, 2023). For the HMGA2 subclass, UL-increased proteins were enriched for transcriptional and ribosome-related functions (**Figure S2C**), which fits prior subtype definitions where HMGA2 activation is coupled to strong transcriptional reprogramming (Mehine *et al*., 2016). The FH subclass showed increased upregulation of proteins involved in metabolic and mitochondrial programs (**Figure S2D**). This aligns with the known metabolic reprogramming of FH-deficient leiomyomas, a consequence of fumarate accumulation that often leads to NRF2 pathway activation and downstream signatures of oxidative and metabolic stress (Mehine *et al*, 2022).

Within this context, PAGE4 emerged again as a prominently UL-increased protein, with the strongest increase observed in the MED12 subclass in our DIA proteome (**Figure 3B**). Together with the subtype-resolved pathway patterns (**Figure S2A**), these results motivated us to test whether PAGE4 enrichment is also reflected at the transcript level. PAGE4 expression indeed was significantly elevated in MED12-mutant ULs compared to normal myometrium and other driver-defined subtypes in an external RNA-seq resource (**Figure 3C**) (Berta *et al*., 2021). This upregulation distinguished MED12-mutant tumors from those driven by HMGA2, YEATS4 (SRCAP complex subunit), OM* (other SRCAP complex subunits), FH, and unknown (lacking defined drivers) tumors. Importantly, the MED12 group in this external dataset comprises multiple MED12 mutation types and is not restricted to the p.G44D variant, indicating that elevated PAGE4 expression is associated broadly with MED12-mutant leiomyomas rather than a single hotspot allele.

### MED12-mutant ULs show coordinated PAGE4 upregulation and site-specific phosphoregulation

Having established concordant PAGE4 elevation at both the protein and mRNA levels, we then asked whether PAGE4 phosphorylation reflects genuine site-specific regulation in MED12-mutant ULs rather than a byproduct of increased PAGE4 abundance. To address this, we normalized phosphosite intensities to their parent proteins using the matched global proteome (protein-adjusted, site-level effects). We then performed subtype-resolved differential testing (UL vs. matched MM) and visualized the results as genotype-specific volcano plots (**Figure 3D**). As in the discovery cohort, MED12 subclass showed strong protein-adjusted increases at PAGE4 T51 and PAGE4 T85(**Figure 3D; Table S2**), supporting the conclusion that PAGE4 hyperphosphorylation represents a genuine phosphorylation in this molecular context rather than a byproduct of elevated PAGE4 protein abundance. Importantly, the expanded cohort also clarified how PAGE4 phosphoregulation behaves across other UL subclasses. In the COL4A5-COL4A6 subclass, PAGE4 T85 also remained among the UL-increased sites after protein adjustment (**Figure 3D**), consistent with these tumors sharing the MED12-mutant background in our cohort and suggesting that the MED12-associated phosphorylation state can extend into this genomic context. In contrast, HMGA2 tumors did not show PAGE4 sites among the significant phosphosite changes (**Figure 3D)**, indicating that PAGE4 phosphoregulation is not a general feature of HMGA2-activated ULs in this dataset. Notably, the FH subclass showed an opposite PAGE4 behavior at the phosphosite level (**Figure 3D; Table S2**). This pattern argues that PAGE4 phosphorylation is not universally “MED12-specific” in the sense of being unchanged elsewhere. Instead, the data support a more precise statement: MED12-driven ULs are characterized by increased PAGE4 phosphorylation at T51 and T85, whereas FH-deficient ULs can display directionally distinct PAGE4 site regulation, pointing to different upstream signaling or phosphoregulation logic in that driver class.

To nominate upstream kinases that may contribute to the MED12 ULs phosphosite program, we performed kinase-substrate scoring on the MED12 phosphoproteome (Kim *et al*, 2021b), generating a combined kinase-substrate score matrix (**Figure 3E**). This analysis indicated preferential activity of CMGC family kinases, including CDKs and MAPKs, consistent with the broader MED12 phosphosite signature. Among individual kinases, HIPK2 emerged as the top candidate predicted to phosphorylate PAGE4 at T51 and T85, both sites fitting their known serine/threonine consensus motifs. Notably, HIPK2 is a stress-responsive kinase previously implicated in regulating oncogenic phosphorylation in hormone-related tumors, particularly breast cancer (Lin *et al*, 2018a), suggesting a potentially conserved mechanism in UL tumorigenesis. Collectively, the DIA validation establishes that PAGE4 is elevated at both the protein and protein-adjusted phosphosite levels specifically in MED12-mutant leiomyomas, setting the stage for the mechanistic experiments that follow.

### Phosphovariant-specific interactomes of PAGE4 and MED12 p.G44D reveal a functional shift toward RNA metabolism and mitochondrial pathway activation in UL

Building on the discovery that PAGE4 is hyperphosphorylated at Thr51 and Thr85 in MED12-mutant ULs (**Figure 3D**), we sought to define the functional consequences of these modifications. We prioritized sites T51 and T85, along with a previously implicated site (S9) (**Figure 4A**), due to their established regulatory potential and significant upregulation in patient tissues (He *et al*, 2015; Kulkarni *et al*, 2017; Mertins *et al*, 2014; Mooney *et al*, 2014; Shiromizu *et al*, 2013; Zhou *et al*, 2013). To systematically decipher the phosphorylation-dependent interactome of PAGE4, we engineered a panel of inducible cell lines expressing either MED12 (WT or p.G44D) or PAGE4 variants. The PAGE4 variants included wild type, phosphomimic to simulate constitutive phosphorylation, and phosphodead to block phosphorylation mutants at these three key residues (**Figure 4A**), enabling comparative interactome mapping across 11 distinct engineered cell lines. Each bait was profiled by affinity purification (AP) and proximity-dependent biotin labeling (PL) mass spectrometry to capture both stable and neighborhood interactions (**Figure 4A**). We identified hundreds of high-confidence interactors (HCIs) after stringent filtering for each bait (**Figure 4B; Table S3**). Notably, single-site mutant (Mut6, T51E) and triple-site mutant (Mut1, S9D/T51E/T85E) of PAGE4 showed reduced stable protein interactions and increased proximity interactions (**Figure 4B**). Among the AP-derived HCIs, the phosphomimic triple mutant (Mut1: S9D/T51E/T85E) retained only 73% of the interactors captured by PAGE4 WT, representing a net loss of approximately 27% of stable associations (**Figure 4B; Table S3**). This reduction was not uniform: while interactions with basal transcription factors and cell-cycle regulators were preferentially lost, associations with Mediator subunits and RNA-processing factors were selectively retained. Conversely, the proximity labeling data showed an expansion of the phosphomimic neighborhood, consistent with a model in which phosphorylation sharpens the stable core of PAGE4 interactions while broadening its transient, functional milieu. This dual behavior supports the interpretation that phosphorylation functions as a specific filter, sculpting the PAGE4 interactome from a broadly engaged state toward a focused, Mediator-centric regulatory module.

**Figure 4:**
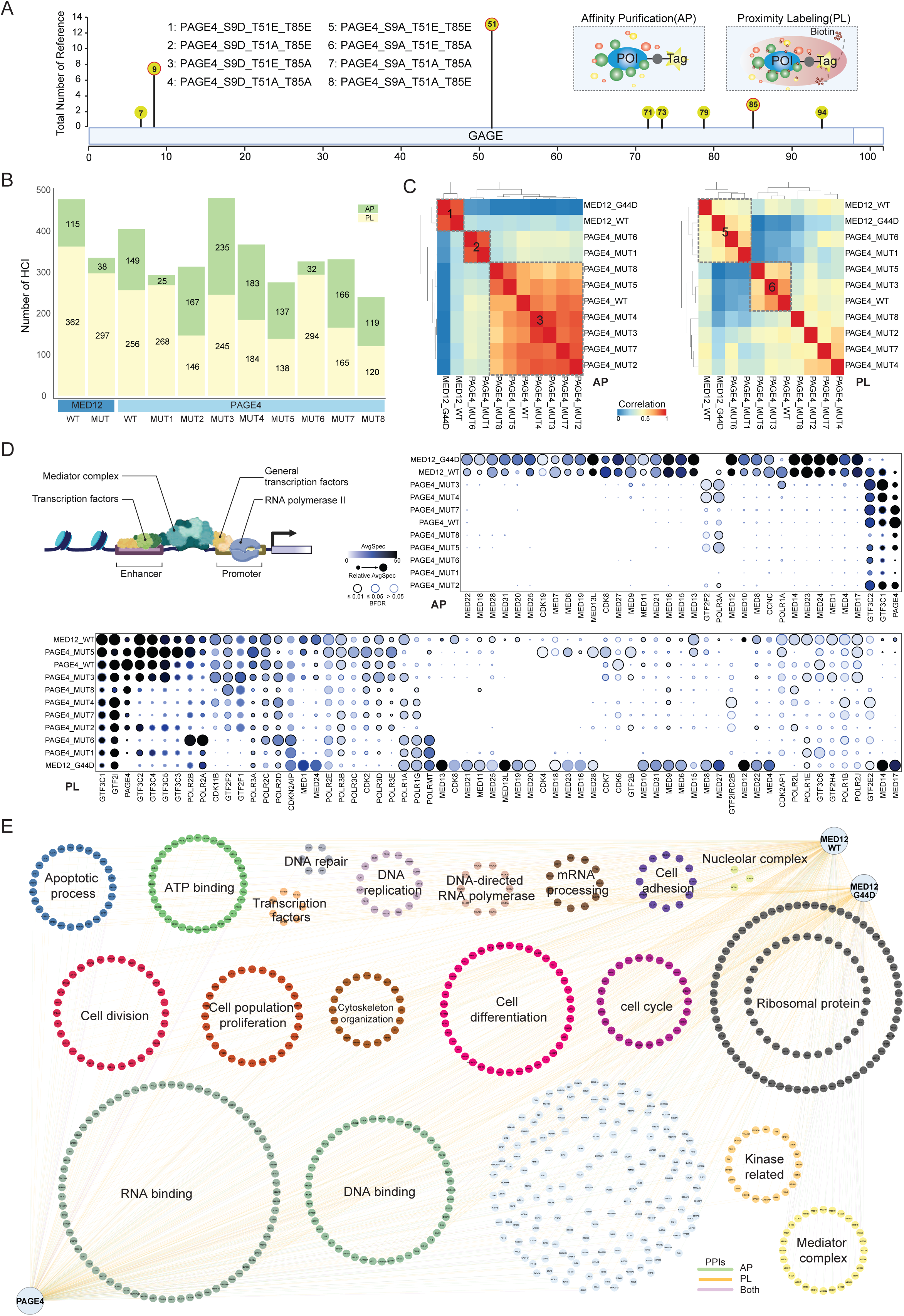
Phosphorylation remodels the PAGE4 interactome toward transcriptional machinery. (**A**) A detailed map of PAGE4 phosphosites is presented alongside a panel highlighting the most frequently researched sites based on literature support counts, with bar height reflecting the number of relevant research articles. The three most extensively studied sites (S9, T51, T85) were selected for functional characterization. Eight PAGE4 variants were generated: wild-type, single-site phosphomimics (S9D, T51E, T85E), single-site phosphodead mutants (S9A, T51A, T85A), and a triple phosphomimic (S9D/T51E/T85E). Each variant was analyzed using complementary approaches: AP-MS for stable interactions and PL-MS with BioID for transient/weak interactions. (**B**) Bar plot showing high-confidence interactors identified per bait after stringent filtering (SAINT BFDR < 0.05, CRAPome < 20%, average spectral-count FC > 3). Stacked bars distinguish interactions captured by AP (light) versus PL (dark). (**C**) Bait-to-bait correlation analysis reveals phosphorylation-dependent interactome reorganization. Heatmaps show Pearson correlation coefficients calculated from interactor spectral count profiles for AP-MS (left) and PL-MS (right). Hierarchical clustering identifies distinct interactome signatures. (**D**) The schematic representation of the transcription machinery illustrates the interaction of Mediator subunits, general transcription factors (GTFs), CDKs/CCNC, and RNA polymerases at the promoter and enhancer regions. The focused interaction matrix follows, displaying dot area encoding averaged spectral signal, fill intensity reflects relative enrichment; the ring shade around each dot indicates SAINT confidence (BFDR ≤0.01, ≤0.05, >0.05). (**E**) A PAGE4-centered PPI map summarizing functionally annotated neighborhoods. Edge color denotes the evidence source: AP (yellow), PL (blue), or both (orange). MED12-WT and MED12-mutant nodes are overlaid to visualize shared interactors with PAGE4.

To determine how phosphorylation changes the PAGE4 protein interaction landscape, we performed comparative analysis of all baits, with interaction profiles clustered by correlation (**Figure 4C)**. In AP data, hierarchical clustering revealed three distinct groups: MED12 baits (Cluster 1), a phospho-favored PAGE4 subgroup (Cluster 2), and the remaining PAGE4 variants (Cluster 3). Notably, Cluster 2 was exclusively populated by the triple phosphomimic Mut1 (S9D/T51E/T85E) and the single mutant Mut6 (T51E). In contrast, the PL data showed the same phosphomimic PAGE4 variants (Mut1 and Mut6) co-clustered with the MED12 baits (Cluster 5), while other PAGE4 forms segregated separately (**Figure 4C**).

We next focused on interactions with core transcriptional machinery, including Mediator subunits, RNA polymerase I/II/III subunits (POLR), general transcription factors (GTF2/GTF3C), and CDKs (CDK7/8/9/11B/19; CCNC). This focused view captures the coupling of Mediator to pre-initiation complex (PIC) assembly, Pol III machinery, and kinase-regulated transcription in the context of MED12, MED12-G44D, PAGE4, and related mutations (**Figure 4D; Table S3**). In AP, the MED12_G44D mutant specifically lost interactions with PAGE4 and POLR3A, while its associations with POLR1A and the GTF3C1/3C2 complex were substantially weakened (**Figure 4D; Table S3**). This indicates that the MED12-G44D mutation disengages the Mediator complex from RNA polymerase III machinery and partially from RNA polymerase I, thereby reducing support for basal transcriptional programs governing tRNA and rRNA synthesis. Strikingly, phosphomimic PAGE4 variants (Mut1: S9D/T51E/T85E; Mut6: T51E) also exhibited specific disengagement from core transcription machinery. Both mutants lost stable interactions with POLR3A and GTF2F2, a general transcription factor that supports Pol II initiation and elongation (**Figure 4D; Table S3**). This pattern aligns with a model in which PAGE4 phosphorylation functions as a molecular switch, redirecting its associations away from stable integration into basal transcription complexes and toward Mediator and other transcriptional co-regulators, as corroborated by subsequent clustering analyses.

The PL data provided a congruent perspective on the local protein environment. Both the MED12-G44D mutant and phosphomimic PAGE4 variants displayed the loss of proximity to key regulators, including CDK7 (the TFIIH kinase responsible for Pol II CTD Ser5 phosphorylation), GTF2B (an essential factor for PIC assembly), and cell cycle-related kinases CDK6 and CDK11B (**Figure 4D; Table S3**). The combined loss of proximity to CDK7 and GTF2B implies a weaker functional link to the early stages of Pol II transcription initiation.

To visualize these relationships, we mapped the interactomes of MED12_WT, MED12-G44D, and PAGE4_WT, grouping preys by function (**Figure 4E**). All three baits interacted with large RNA-binding and ribosomal protein clusters and many Mediator subunits, indicating a common scaffold that links them to transcriptional regulation and RNA handling. Specifically, the MED12 wild type shows densely connecting to nucleolar/Pol I-III and PIC/Pol II GTF circles, consistent with coupling to rRNA/tRNA synthesis and early Pol II initiation (Svejstrup *et al*, 1997). In contrast, the G44D displayed sparser connections into the nucleolar/Pol I-III cluster and fewer ties to PIC/Pol II GTFs, indicating reduced support for housekeeping transcription. PAGE4 connected strongly to mediator, RNA-binding, and ribosomal circles, with fewer direct links into Pol I/III and PIC nodes (**Figure 4E**). This matches the phospho-switch model where PAGE4 shifts from basal machinery toward Mediator-centric regulation.

Collectively, our data reveals a coherent model that the MED12-G44D mutation and PAGE4 phosphorylation drive transcriptome reallocation by uncoupling mediator and its associated factors from the basal transcription machinery for Pol I, II, and III. The shift from global housekeeping transcription toward more selective, stress-adaptive programs aligns with observed upstream regulator phenotypes involving extracellular matrix remodeling(Winkler *et al*, 2020), altered cellular mechanics, and RNA metabolism (Matia-González *et al*, 2021).

Building on our discovery that phosphomimics PAGE4 variants Mut1 (S9D/T51E/T85E), Mut6 (T51E), and MED12-G44D converge on a similar disruption of the basal transcription machinery (**Figure 4D**), we sought to define the specific cellular programs to which their interactions are redirected.

A comparative analysis of interactors revealed that the PAGE4 phosphomimic Mut1 and Mut6 gained associations with a distinct set of proteins involved in RNA handling (**Figure 5A; Table S3**), such as ribosomal proteins RPL37A and RPL27(Brodersen & Nissen, 2005), and factors controlling protein homeostasis and vesicular trafficking, including CKMT1A, GPATCH4, LGALS3BP and SLC25A10 (Pathan *et al*, 2019). In the MED12 p.G44D dataset, the bait (MED12-G44D) itself is strongly reduced compared with MED12 WT, consistent with decreased recovery of the mutant complex. Despite this global decrease in bait abundance, the volcano plot shows a selective loss of a subset of interactors rather than a uniform collapse of all prey signals. Specifically, many ribosomal proteins display marked decreases, indicating that the interaction changes are not solely explained by reduced bait amount but reflect differential binding/association in the mutant background (**Figure 5A; Table S3**).

**Figure 5:**
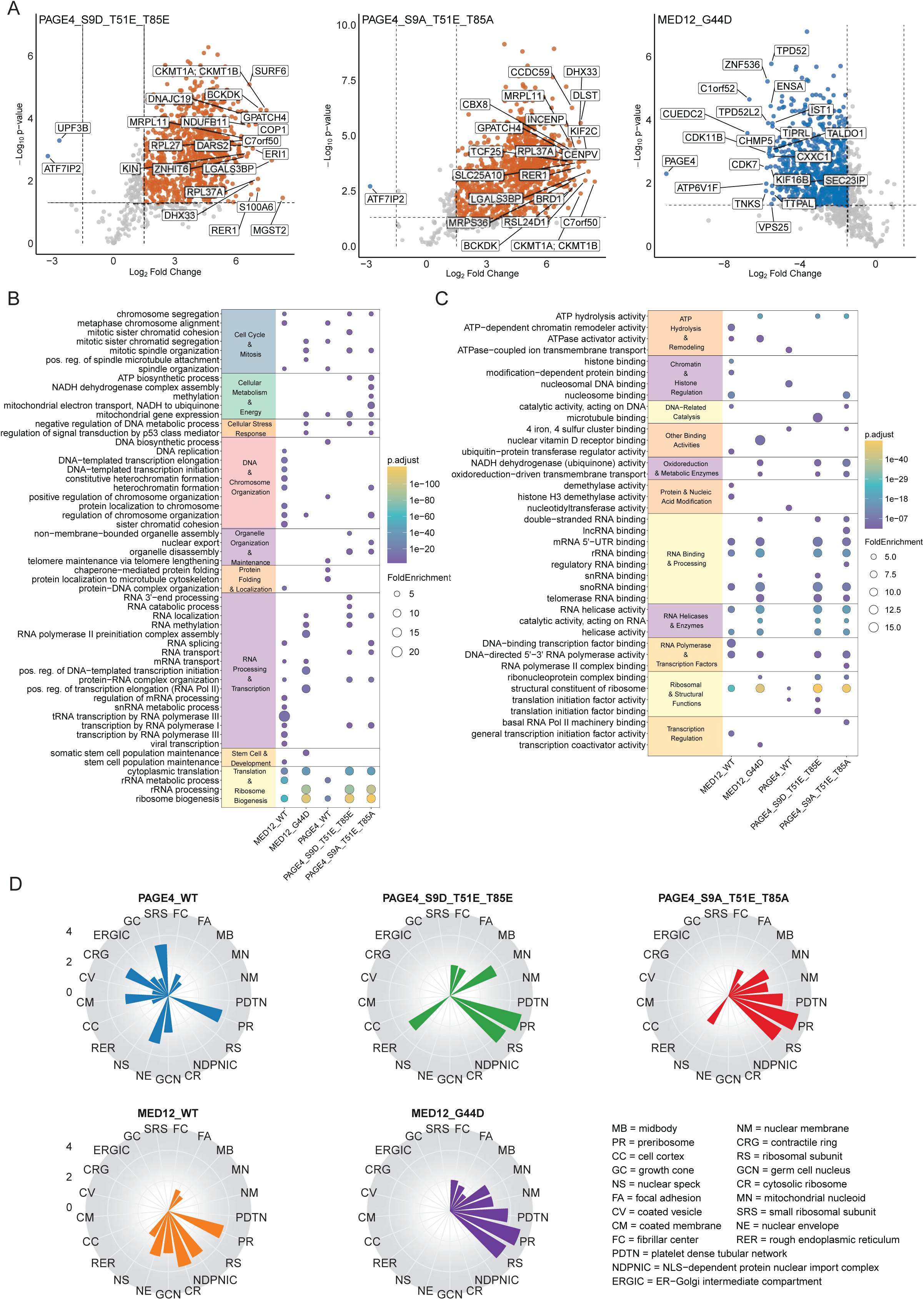
Phosphorylation-dependent functional remodeling of PAGE4 and MED12 interactomes. (**A**) Volcano plot of differentially interacting proteins, comparing PAGE4 single-site and triple-site phosphomutants, and MED12 G44D-mutant to their respective wild-type counterparts. Significantly enriched interactors are highlighted (|log₂FC| > 1.5, p < 0.05). (**B)** Gene Ontology (GO) Biological Process (BP) enrichment analysis of interactors, illustrating functional shifts in PAGE4 and MED12 mutants. (**C)** GO Molecular Function (MF) enrichment analysis, showing functional category enrichment for mutant and wild-type PAGE4 and MED12 interactors. (**D)** GO Cellular Component (CC) enrichment analysis, demonstrating distinct subcellular localizations of interactors for PAGE4 WT, phosphomutants, MED12 WT, and MED12 G44D, emphasizing phosphorylation-dependent changes in protein complex organization.

Among the most notable phosphorylation-dependent interactors, the estrogen receptor coactivator PELP1 (proline-, glutamic acid-, and leucine-rich protein 1) was identified exclusively in the phosphomimic PAGE4 triple mutant (Mut1: S9D/T51E/T85E) by PL (Table S3). Critically, PELP1 was absent from all other PAGE4 variants and from both MED12 WT and MED12 G44D interactomes, indicating that its recruitment is strictly contingent on multisite PAGE4 phosphorylation. PELP1 is a well-established scaffolding protein that couples estrogen receptor signaling to chromatin remodeling and has been implicated in hormone-dependent tumor progression in breast and ovarian cancers. Its phosphorylation-dependent association with PAGE4 in this context suggests a mechanism through which hyperphosphorylated PAGE4 may directly interface with estrogen signaling pathways in MED12-mutant uterine leiomyomas.

This reprogramming of the PAGE4 interactome was reflected in its functional associations. Upon phosphorylation, PAGE4-associated proteins became significantly enriched for roles in RNA processing, ribosome biogenesis, and translation, alongside functions in protein folding and organelle transport (**Figure 5B and 5C**). Concurrently, its connections to cell-cycle progression, DNA binding, and chromatin organization were markedly reduced. The MED12-G44D mutant again displayed a congruent profile, with diminished representation of factors involved in core transcription.

The subcellular context of this shift was clear: the interaction network of phosphomimic PAGE4 relocated from chromosomal and transcription initiation compartments toward ribosomal, nucleolar, and secretory organelle environments (**Figure 5D**). This spatial redistribution aligns with its newfound affinity for RNA and protein maintenance machinery.

Together, these data demonstrate that phosphorylation at T51, alone or in combination with S9 and T85, reprograms PAGE4 from a factor associated with basal transcription to one engaged with post-transcriptional RNA handling and proteostasis. This functional shift mirrors the reallocation of transcriptional resources observed with the MED12-G44D mutation, reinforcing a model where both perturbations converge to divert cellular output from housekeeping functions toward specialized, stress-adaptive programs.

### Divergent pathway perturbations define MED12 and PAGE4 functional landscapes across wild-types and variants

To assess the functional impact of PAGE4 and MED12 variants, we performed Reactome pathway enrichment on bait-specific interactors. (**Figure S3**). MED12 WT exhibited strong transcriptional enrichment, with pathways including RNA polymerase I/II/III transcription, chromatin remodeling, and transcriptional regulation by TP53 and E2F6, reinforcing its role in transcriptional control and chromatin dynamics. DNA repair and genome stability pathways were significantly enriched, highlighting MED12 WT’s role in cell proliferation and genome maintenance. The MED12 G44D mutation exhibited reduced enrichment in core RNA polymerase transcription pathways, yet showed a shift toward transcriptional regulation of white adipocyte differentiation, MECP2-regulated transcription, and SUMOylation-related processes, suggesting altered transcriptional priorities. DNA repair pathway enrichment was less pronounced compared to the WT, with a greater emphasis on mitochondrial RNA metabolism and transport, suggesting plausible reprogramming of RNA processing in the mutant state (**Figure S3**).

In contrast, PAGE4 WT displayed a distinct enrichment for oncogenic signaling and cytoskeletal regulation, with significant pathways including Rho GTPase signaling, vesicle-mediated transport, and Golgi-to-ER retrograde trafficking (**Figure S3**), reinforcing its potential role in modulating intracellular architecture and signaling networks. FGFR2-mediated signaling pathways were prominently enriched, along with SUMOylation, apoptosis regulation, and metabolic processes such as PPARA-activated gene expression and glycogen metabolism, linking PAGE4 to metabolic adaptation and stress response in the tumor microenvironment. The single-site mutant (Mut6) exhibited enhanced enrichment in RNA transport and metabolism pathways, indicating a potential influence on RNA stability and cellular energy regulation. In contrast, the triple-site mutant (Mut1: S9D/T51E/T85E) showed a shift toward DNA repair regulation, with significant enrichment in base excision repair, translation synthesis, and SUMOylation of DNA damage response proteins, as well as a greater association with mitotic regulation and oxidative metabolism pathways, suggesting a functional divergence from the wild-type protein. Collectively, these findings highlight that MED12 and PAGE4 variants exhibit distinct yet overlapping functional signatures, with MED12 prioritizing transcription and genome stability, while PAGE4 integrates oncogenic signaling and cytoskeletal regulation. Variants display divergent functional shifts, with MED12 G44D showing transcriptional and repair deficits, PAGE4 single-site mutant (Mut6) acquiring enhanced RNA transport activity, and triple-site (Mut1: S9D/T51E/T85E) exhibiting DNA repair and metabolic regulatory features, underscoring the functional plasticity of these interactomes.

### Phosphorylation of PAGE4 and MED12 G44D mutation influence transcriptional and signaling pathways in UL

To evaluate the functional impact of MED12 mutations and PAGE4 phosphorylation on key signaling pathways, we conducted luciferase reporter assay across PAGE4 and MED12 variants (**Figure S4A-S4F**). As reported for UL pathobiology, the MAPK/ERK (**Figure S4C**) and estrogen signaling (**Figure S4E**) pathways are typically hyperactivated in UL, contributing to proliferation and extracellular matrix remodeling (Borahay *et al*, 2017). However, PAGE4 phosphovariants including Mut1 (S9D/T51E/T85E) and Mut6 (T51E), exhibited reduced ERK and estrogen reporter activity relative to PAGE4 wild type (**Figure S4**). Rather than directly inhibiting MAPK/ERK and estrogen signaling, phosphorylated PAGE4 (Mut1 and Mut6) may engage alternative functional networks, potentially favoring RNA metabolism and mitochondrial processes, as suggested by interactomics analyses (**Figure 5B**). This shift may reflect altered interactions with transcriptional co-factors required for ER-mediated gene expression and MAPK activation, potentially affecting UL-associated signaling. The suppression of Wnt signaling in the triple-site mutant PAGE4 (Mut1: S9D/T51E/T85E) (**Figure S4D**) further raises the possibility of a functional change, as Wnt activation is commonly implicated in UL pathogenesis(El Sabeh *et al*., 2021). Conversely, increased activation of the cell cycle/pRb-E2F pathway (**Figure S4A**) in PAGE4 phosphovariants may indicate an altered role in cell cycle regulation, a possibility consistent with PAGE4’s known involvement in androgen receptor signaling in prostate cancer. (Obinata *et al*, 2024)

For MED12 G44D, reduced MAPK/ERK and Wnt signaling activity (**Figure S4**) suggests potential disruptions in transcriptional regulation that may be relevant to uterine smooth muscle homeostasis. Given the high prevalence of MED12 mutations in ULs, these findings are consistent with the hypothesis that the G44D variant could contribute to changes in transcriptional programs and extracellular matrix remodeling, which are central to tumor development. (Moyo *et al*., 2020) The mutation’s association with altered RNA polymerase II-mediated transcription may also reflect broader shifts in gene regulatory networks that could influence the epigenetic landscape of UL.

Together, these findings suggest that phosphorylation-dependent PAGE4 modulation and MED12 G44D-driven transcriptional alterations intersect with core pathways involved in cell cycle progression, proliferation, and cellular stress responses.

### Transcriptional landscape of MED12 G44D mutation and PAGE4 phosphovariants reveals regulatory shifts in RNA polymerase II (POLRII) recruitment and transcription factor binding

Building on the pathway reporter assays that indicated distinct transcriptional outputs for phosphorylated PAGE4 and MED12 G44D, we asked whether these functional differences were reflected in altered chromatin association and transcription factor motif usage. ChIP-seq analysis show that both MED12 and PAGE4 wild type exhibit sharp, symmetric peaks centered on transcription start sites (**Figure 6A; Figure S5A**). In contrast, MED12 G44D and the phosphorylated PAGE4 (Mut1 and Mut6) display slightly broader or lower peaks (**Figure 6A; Figure S5A**), suggesting altered binding distribution or reduced occupancy near active promoters (**Figure S5B**). HOMER motif analysis revealed overlapping but distinct motif repertoires between wild-type and mutant proteins (**Figure 6B**). We observed that MED12 WT supports transcriptional maintenance and differentiation, enriched for AP-1, E2F3, and KLF factors. In contrast, the G44D mutant exhibited a stress-adaptive signature, marked by increased occupancy of ETS family factors (ELF1, ETS1, ETV2) and nuclear receptors (FOXA1, FOXM1, THRα), consistent with TF programs linked to proliferation, (ETS1), invasion (ETV2), and hormone-independent growth (FOXA1) (Dhara *et al*, 2022; Luk *et al*, 2018). Similarly, PAGE4 WT maintained AP-1-driven transcription, whereas phosphorylation induced an expanded motif repertoire. Triple-site phosphorylation on PAGE4 (Mut1: S9D/T51E/T85E) showed enrichment for FOXA1, ETV2, and HLF, suggesting activation of transcriptional programs linked to developmental regulation and plasticity, while the single-site mutant (Mut6) retained WT-like features but gained TEAD and CEBP:AP1 motifs (**Figure 6B**), suggesting progressive transcriptional rewiring(Currey *et al*, 2021; Han *et al*, 2023; Kim *et al*, 2023; Toh *et al*, 2017; Wang *et al*, 2024).

**Figure 6:**
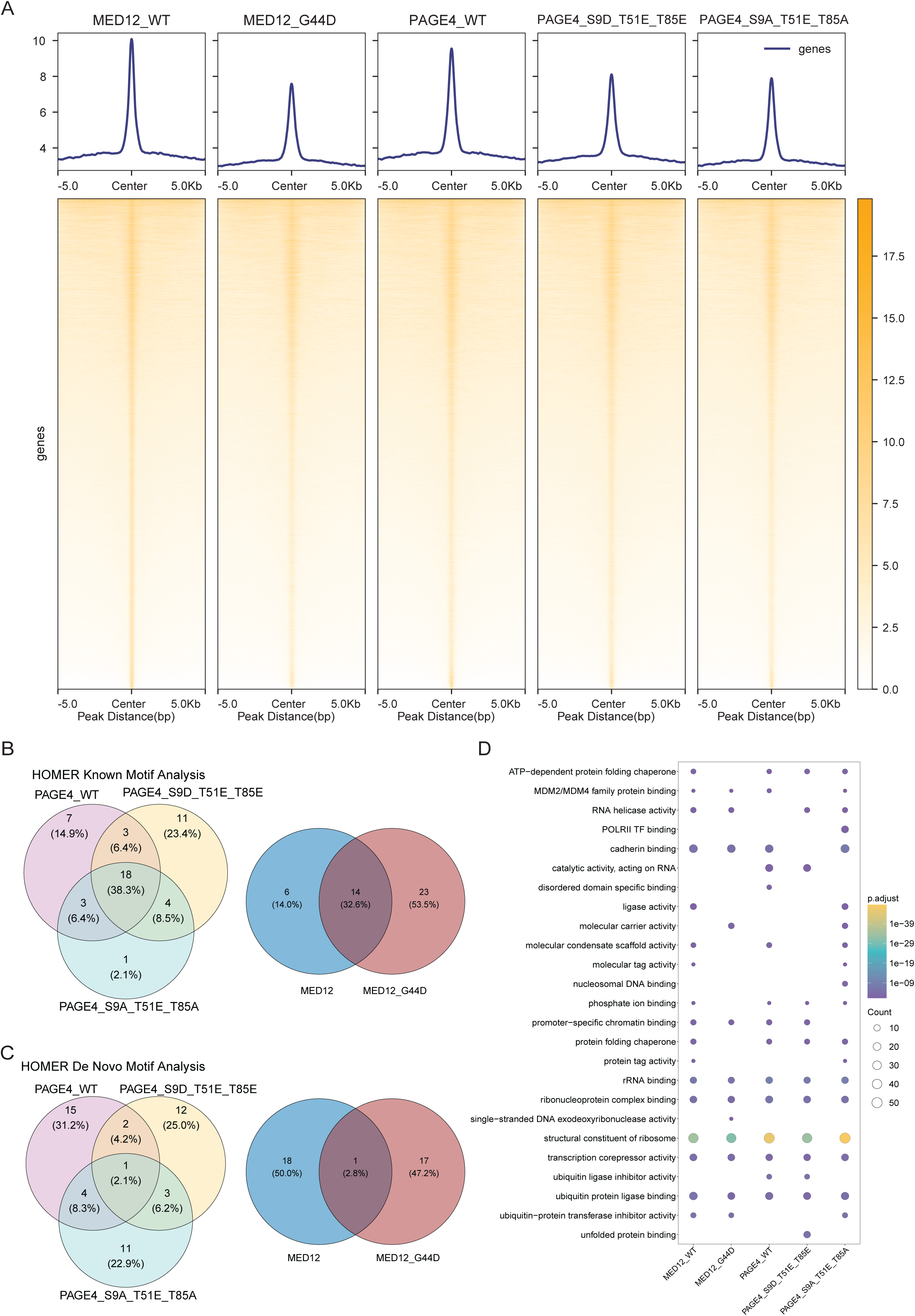
MED12 G44D mutation and PAGE4 phosphorylation drive convergent transcriptional reprogramming in uterine leiomyomas. **(A)** Genome-wide POLII occupancy profiling by ChIP-seq reveals phosphorylation-dependent transcriptional rewiring. Stable cell lines with MAC3-tagged constructs (MED12 WT, MED12 G44D, PAGE4 WT, or PAGE4 phosphomimic mutants), followed by chromatin immunoprecipitation using anti-POLII antibody and high-throughput sequencing. Heatmaps display POLII signal intensity (±3 kb from transcription starts sites, TSS) across all detected genes, hierarchically clustered by binding pattern. Bar plots (right) show the number of POLII-enriched peaks per condition (fold-enrichment > 2, FDR < 0.05). Color scale represents normalized read density (RPKM). (**B)** HOMER known motif analysis reveals distinct transcription factor binding patterns, with MED12 WT enriched for transcriptional maintenance factors (AP-1, E2F3, KLF), while MED12 G44D exhibits increased binding of stress-adaptive ETS factors (ELF1, ETS1, Etv2) and nuclear receptors (FOXA1, FOXM1, THRa). PAGE4 WT maintains AP-1-driven transcription, whereas phosphorylation induces a shift toward FOXA1, ETV2, and HLF motifs, promoting a stem-like transcriptional state. (**C)** HOMER de novo motif analysis demonstrates MED12 G44D-driven loss of differentiation-associated motifs (MEF2C, PITX2, Nkx2.1) and gain of stress regulators (ATF3, IRF4, p53). PAGE4 phosphorylation disrupts cell cycle-associated transcription factors (FOSL2, JUND), leading to enrichment of epigenetic regulators (ZNF384, Six3, ZBTB6), while ZNF135 remains a conserved regulatory signature across PAGE4 conditions. (**D)** Functional annotation of POLII-bound genomic regions reveals distinct molecular function profiles. Genes associated with POLII ChIP-seq peaks were subjected to GO molecular function enrichment analysis to identify functional categories of transcriptional targets. Bar plots display significantly enriched GO terms (FDR < 0.05, fold-enrichment > 2) for each condition. Enrichment analysis performed using DAVID and Metascape tools.

HOMER *de novo* analysis (**Figure 6C; Table S4**) further revealed MED12 G44D-driven reduced enrichment of differentiation-associated motifs (MEF2C, PITX2, Nkx2.1) and increased association with stress-related regulators (ATF3, IRF4, p53), shifting MED12’s role toward cellular stress adaptation (**Figure 6C; Table S4**). Meanwhile, PAGE4 phosphorylation disrupted core transcriptional control, replacing cell cycle-associated factors (FOSL2, JUND) with ZNF384, Six3, and ZBTB6 (**Figure 6C**), suggesting alternative epigenetic regulation. (Shetty *et al*, 2022) The consistent presence of ZNF135 across all PAGE4 conditions highlights a conserved regulatory signature. Molecular function (MF) analysis of ChIP-seq peak annotations reinforced findings from *de novo* and known motif analyses (**Figure 6D; Table S4**) supported these shifts, with MED12 WT enriched for chromatin remodeling and transcriptional regulatory functions, G44D enriched for stress-associated functions, and phosphorylated PAGE4 mutants enriched for nucleosomal DNA binding and ribosomal interaction terms. This post-transcriptional reprogramming was further reinforced by the phospho-PAGE4 interactome, which prominently recruited DEAD-box RNA helicases and ribosomal proteins. DDX21 emerged as a major phosphorylation-dependent interactor in the triple phosphomimic PAGE4 (Mut1: S9D/T51E/T85E; Table S3). In parallel, 254 unique ribosomal proteins were detected across phosphomimic PAGE4 variants, including 87 specific to the triple phosphomimic (**Table S3**). Combined with the concurrent loss of stable interactions with basal transcription factors (**Figure 4D**), these data support a trajectory in which PAGE4 phosphorylation shifts function from promoter-proximal transcriptional roles toward RNA processing and translational control. Consistent with this model, MED12-mutant leiomyoma proteomes showed upregulation of spliceosome and SRP-dependent translational pathways (**Figure 1E**), suggesting that the phospho-PAGE4 interactome may act as a molecular node organizing tumor-level gains in RNA handling and translation.

## Discussion

ULs are highly fibrotic tumors characterized by excessive ECM deposition, altered mechanical signaling, and distinct transcriptional programs (Buyukcelebi *et al*, 2023b; Koohestani *et al*, 2013; Rogers *et al*, 2008). By integrating proteomics, phosphoproteomics, AP and PL interactomics, reporter assays, RNA-seq, and ChIP-seq, we support a model in which MED12 mutation and phosphorylation-dependent regulation of the intrinsically disordered protein PAGE4 converge on fibrogenesis, stress adaptation, and transcriptional reprogramming. Consistent with the fibrotic character of ULs, our proteomic comparisons highlighted increased abundance of multiple extracellular matrix and matrix-remodeling factors, including collagens (COL1A1, COL3A1, and COL4A1) and ECM-associated proteins such as POSTN, together with enzymes linked to matrix remodeling and fibrotic signaling such as MMP2, TGFB3, and PLOD1. In parallel, functional enrichment of UL-altered proteins and phosphosites pointed to coordinated changes in ECM-receptor interaction, integrin-linked adhesion/focal-adhesion modules, and regulation of the actin cytoskeleton, supporting altered cell-matrix coupling and structural remodeling as recurring features of the UL state. These features are consistent with a mechanically altered UL state in which integrin-linked signaling and cytoskeletal remodeling accompany ECM expansion.

PAGE4 emerged as a consistently induced marker in MED12-mutant ULs at protein, transcript, and IHC levels, with cytoplasmic-predominant staining that cleanly distinguishes UL from matched myometrium. This extends prior observations suggesting PAGE4 expression in UL tissues while providing multi-layer evidence for its enrichment in the MED12-mutant context (Fu *et al*, 2020). Its dual nuclear-cytoplasmic localization and consistent induction further support PAGE4 as a robust diagnostic marker (Jakimiuk *et al*., 2004; Leitao *et al*., 2004; Mehasseb *et al*., 2011). Crucially, DIA phosphoproteomics revealed robust, protein-adjusted increases at PAGE4-T51 and -T85 specifically in MED12-mutant tumors, expanding its known functional landscape beyond prostate cancer (Fu *et al*., 2020; Kulkarni *et al*., 2017; Lin *et al*., 2018a; Mooney *et al*., 2014). Notably, PAGE4 Thr51 showed the opposite direction in FH-deficient tumors (decreased), indicating that PAGE4 phosphorylation is driver-context dependent rather than universally “on” across UL classes. Thus, the coordinated increase of Thr51 and Thr85, together with elevated PAGE4 abundance, is characteristic of the MED12-mutant background. Kinase-substrate inference pointed to CMGC family activity. Within this group, HIPK2 was the top candidate for PAGE4-T51/T85, suggesting stress-responsive control of PAGE4 in UL (Chowdhury *et al*, 2023; Kim *et al*, 2021a). Together, these data link stress-adaptive kinase activity to PAGE4 regulation in UL and align with the view of IDPs as phosphorylation-gated molecular rheostats in hormonally responsive tissues (He *et al*., 2015; Kulkarni *et al*., 2017; Lin *et al*., 2018a).

Functionally, phosphomimic PAGE4 variants (Mut1: S9D/T51E/T85E; Mut6: T51E) remodeled the interactome away from pre-initiation and Pol I/III machinery toward RNA handling, ribosome biogenesis, proteostasis and organelle traffic. These findings reinforce a role for PAGE4 in metabolic adaptation and cytoskeletal reorganization, which are key processes associated with tumor evolution(Ning *et al*, 2024; Wang *et al*, 2022; Yang & Al-Hendy, 2023). PAGE4 phosphorylation broadly rewired its network, reducing stable interactors by 27% while expanding proximity associations, consistent with conversion from a broadly connected hub to a more selective signaling module. This specificity-filter model has precedent in intrinsically disordered proteins (IDPs), where phosphorylation can induce disorder-to-order transitions that collapse the conformational ensemble and favor defined binding interfaces (Newcombe *et al*, 2022). In UL, this selectivity toward Mediator-associated and RNA-processing complexes over basal transcription machinery suggests that phospho-PAGE4 acts at the interface of transcription and translation.Similarly, MED12 G44D mutation altered the interactome landscape, suppressing wild-type transcriptional programs in favor of stress-adaptive and RNA-centric pathways. Both PAGE4 phosphovariants and MED12 G44D showed decreased MAPK/ERK and Wnt signaling activities, (Borahay *et al*., 2017; El Sabeh *et al*., 2021; Obinata *et al*., 2024) even though these pathways are typically hyperactivated in UL, suggesting that these molecular changes may reflect a decoupling of classical proliferative signals in favor of alternative adaptation strategies.

Transcriptome-wide analysis of POLRII occupancy and motif enrichment revealed further evidence of reprogrammed gene regulation. MED12 G44D favored binding to stress-associated motifs such as ATF3, IRF4, and p53 (Dhara *et al*., 2022; Luk *et al*., 2018), while the PAGE4 triple-site variant (Mut1: S9D/T51E/T85E) gained accessibility to developmental and stem-like motifs including FOXA1, ETV2, and HLF (Currey *et al*., 2021; Han *et al*., 2023; Kim *et al*., 2023; Toh *et al*., 2017; Wang *et al*., 2024). Consistent with these motif shifts, known motif analyses and GO enrichment converge on altered engagement of the transcriptional machinery, including chromatin remodeling, nucleosomal DNA binding, and ribosomal activity (Shetty *et al*., 2022). KLF family motif enrichment in the phospho-PAGE4 cistrome provides a transcriptional route to smooth muscle de-differentiation, with prominent KLF3 and KLF14 motifs suggesting activation of KLF-driven programs and co-enriched CTCF motifs supporting execution within insulated chromatin domains.Together with the ECM dysregulation, cytoskeletal remodeling, and phosphorylation-mediated interactome shifts mapped here, these findings point to actionable therapeutic opportunities, including targeting CMGC kinases such as HIPK2 or modulating integrin and ECM signaling to disrupt both fibrotic and transcriptional components of UL progression (Borahay *et al*., 2017; Ciebiera *et al*., 2024; Li *et al*, 2021), and they position PAGE4 as a potential dual biomarker and functional phospho-switch readout for therapeutic intervention given its consistently high expression in UL.

Perhaps one of the most integrative findings from our interactome analysis is the loss of PAGE4 binding by MED12 G44D, which suggests a loss-of-restraint mechanism rather than a simple gain-of-function model for the recurrent MED12 mutation. In this framework, WT MED12 normally sequesters PAGE4 within the Mediator complex, integrating it into transcriptional programs that support orderly myometrial function. The G44D mutation, by disrupting this interaction, releases PAGE4 from Mediator-based scaffolding, allowing hyperphosphorylated PAGE4 to redistribute toward the alternative interaction networks we have characterized, RNA processing and ribosome biogenesis complexes, chromatin remodelers (MORC2, H2A variants), and estrogen signaling components (PELP1). PELP1, detected only with fully phosphomimetic PAGE4, offers a direct mechanistic bridge to estrogen dependence by coupling ERα to chromatin-modifying complexes, suggesting that PAGE4 phosphorylation may amplify or redirect estrogen-responsive transcription in MED12-mutant ULs. In parallel, enrichment of H2A variants and MORC2 supports a chromatin-level model in which phospho-PAGE4 associates with nucleosome dynamics (H2A variant-rich regulatory regions) while also engaging silencing machinery (MORC2), consistent with selective activation of UL programs alongside repression of smooth muscle contractility modules. This model elegantly reconciles several observations: (i) the selective association of PAGE4 hyperphosphorylation with MED12-mutant but not other UL subtypes; (ii) the convergence of MED12 G44D and phosphomimic PAGE4 on similar functional outputs (reduced basal transcription, enhanced RNA metabolism); and (iii) the specific disruption of Pol III/Pol I machinery seen in both contexts. Under this model, the pathogenic sequence in MED12-mutant leiomyomas proceeds from somatic mutation (MED12 G44D) to structural uncoupling (loss of MED12–PAGE4 interaction), enabling phosphorylation-dependent rewiring of PAGE4 toward tumor-promoting interaction networks, ultimately driving the transcriptional and phenotypic reprogramming that characterizes these tumors. Future studies should test whether PELP1 depletion, or pharmacologic disruption of the PELP1–PAGE4 interface, alters estrogen-dependent gene expression and proliferation in UL-derived models, and whether perturbing H2A-variant exchange or MORC2 activity modifies the balance between activated (ECM, RNA processing) and repressed (smooth muscle contractility, cytoskeletal organization) programs.

This study has several limitations. First, although our DIA cohort captures several major UL molecular contexts (MED12, HMGA2, FH, and COL4A5-COL4A6), subgroup sizes were unequal, with smaller numbers in the FH and COL4A5-COL4A6 subclasses, which may limit statistical power for subtype-specific conclusions and increase sensitivity to outliers and larger multi-center cohorts would enhance the generalizability of our findings (Galindo *et al*, 2018; Wise & Laughlin-Tommaso, 2016). Second, because all COL4A5-COL4A6 tumors in our cohort also carried concurrent MED12 mutations, we cannot fully separate effects attributable to the COL4A5-COL4A6 deletion from those driven by the MED12-mutant background. Third, kinase activities inferred from phosphoproteomic patterns (e.g., HIPK2) remain predictive and require direct validation using targeted kinase assays and genetic perturbation. Finally, mechanistic and translational conclusions would be strengthened by validating PAGE4 regulation and function in primary patient-derived cells and *in vivo* models, including testing whether modulating candidate kinases or fibrotic pathways changes PAGE4 phosphorylation states and UL-associated phenotypes.

In summary, we uncovered a mechanistic framework that links post-translational regulation and genome remodeling to UL pathogenesis. Our results suggest that MED12 mutation and PAGE4 phosphoregulation contribute to the reprogramming of transcriptional networks, ECM remodeling, and the emergence of a fibrotic, stress-adaptive tumor state in uterine leiomyomas. Together, these findings highlight converging regulatory axes that may be amenable to therapeutic intervention. Future efforts focused on validating kinase-substrate relationships and testing targeted therapies, such as kinase inhibitors or anti-fibrotic agents, could lead to precision-based, minimally invasive treatment strategies for this highly prevalent tumor.

## Material and Methods

### Ethics statement and patient cohort

The study was approved by the Ministry of Social Affairs and Health and the Finnish Institute for Health and Welfare (STM/53/07/2000; THL/1071/5.05.00/2011; THL/151/5.05.00/2017; THL/723/5.05.00/2018; THL/1300/5.05.00/2019; THL/1849/14.06.00/2024), as well as by the Ethics Committee of the Hospital District of Helsinki and Uusimaa (133/E8/03; 408/13/03/03/2009; 177/13/03/03/2016; HUS/2509/2016). All patient samples were collected with written informed consent and pseudonymized to ensure participant confidentiality. The study was conducted in accordance with the Declaration of Helsinki.

This retrospective study utilized archived samples from the Finland Myoma Study cohort, which was previously described and analyzed in detail (Berta *et al*., 2021). The same patient cohort has been reported earlier; however, the present study represents a new and independent proteomic investigation of a selected subset of these samples. Tumors were classified into molecular subtypes based on their driver gene alterations, as previously defined. For the current analyses, matched uterine leiomyoma (UL)-myometrium (MM) pairs were selected to represent the major molecular subtypes: MED12-mutant (n = 9 pairs for discovery DDA and validation DIA), COL4A5-COL4A6 deletion-associated (n = 5 pairs for validation DIA), HMGA2-activated (n = 10 pairs for validation DIA), and FH-deficient (n = 5 pairs for validation DIA). In addition, n = 9 matched pairs were included for immunohistochemistry (IHC) analyses. Detailed sample identifiers, molecular classifications, and their allocation to the discovery (DDA), validation (DIA), and IHC cohorts are provided in the supplementary file.

### Global proteome and phosphoproteome analysis

#### Sample preparation

A total of 20 mg of tissue (dry weight) were lysed in 8 M urea (Sigma-Aldrich). Extracted proteins were reduced with 5 mM Tris(2-carboxyethyl)phosphine (TCEP; Sigma-Aldrich) for 20 minutes at 37°C, followed by alkylation with 10 mM iodoacetamide (IAA; Sigma-Aldrich) for 20 minutes at room temperature in the dark. The urea concentration was diluted by adding 600 µL of ammonium bicarbonate (AMBIC; Sigma-Aldrich) prior to trypsin digestion. Sequencing-grade modified trypsin (Promega) was added at a 1:100 enzyme-to-substrate ratio, and the samples were incubated overnight at 37°C. After digestion, the samples were desalted using C18 macrospin columns (Higgins Analytical).

Phosphopeptide enrichment was performed using titanium (IV) ion immobilized metal ion affinity chromatography (Ti4+-IMAC). The IMAC material was prepared as previously described. (Zhou *et al*., 2013) For enrichment, Ti4+-IMAC beads were loaded onto GELoader tips (Thermo Fisher Scientific) and conditioned with 50 µL of conditioning buffer (50% acetonitrile, 6% trifluoroacetic acid [TFA]) by centrifugation at 150 g. Protein digests were dissolved in loading buffer (80% acetonitrile, 6% TFA) and passed through the spin tips at 150 g. The columns were washed sequentially with 50 µL of wash buffer 1 (50% acetonitrile, 0.5% TFA, 200 mM NaCl) and 50 µL of wash buffer 2 (50% acetonitrile, 0.1% TFA). Bound phosphopeptides were eluted using 10% ammonia, followed by a second elution with 80% acetonitrile and 2% formic acid. The eluted samples were dried in a vacuum centrifuge and reconstituted in 15 µL of 0.1% TFA and 1% acetonitrile.

#### Liquid chromatography-mass spectrometry (LC-MS)

Phosphopeptides were analyzed using a Q-Exactive mass spectrometer (Thermo Fisher Scientific) coupled with an EASY-nLC 1000 system and controlled by Xcalibur software (version 3.0.63). Peptides were separated on a C18 precolumn (Acclaim PepMap 100, 75 µm × 2 cm, 3 µm, 100 Å) and an analytical column (Acclaim PepMap RSLC, 75 µm × 15 cm, 2 µm, 100 Å) using a 120-minute linear gradient with buffer A (0.1% formic acid in 98% HPLC-grade water and 2% acetonitrile) and buffer B (0.1% formic acid in 98% acetonitrile and 2% water).

Data acquisition was performed in positive ion mode, with a full scan range of 200-2000 m/z at a resolution of 70,000, followed by top10 CID-MS2 scans at a resolution of 17,500 and a dynamic exclusion duration of 30 seconds. The resulting MS2 spectral data files (Thermo.RAW) were processed using MaxQuant (version 1.6.0.16; Cox & Mann, 2008) and searched against the human UniProtKB database using the Andromeda search engine (Cox et al., 2011). Static modifications included carbamidomethylation (+57.021 Da) of cysteine, while dynamic modifications included phosphorylation (+79.966 Da) of serine, threonine, and tyrosine, and oxidation (+15.994 Da) of methionine. Precursor and fragment mass tolerances were set to <20 ppm and 0.1 Da, respectively, with a maximum of two missed cleavages allowed. Results were filtered to a false discovery rate (FDR) of <0.05. Phosphorylation sites were further filtered based on a localization probability cutoff of 0.75, and any phosphotyrosine sites identified in control experiments without added kinase were excluded.

#### GO enrichment and statistical analysis

Differentially expressed proteins were identified using log2 fold-change (log2FC) analysis and paired t-tests with an FDR-adjusted significance threshold (*p* ≤ 0.05). Gene Ontology (GO) enrichment analysis was performed using clusterProfiler (Yu *et al*, 2012), with redundant GO terms refined using rrvgo (Sayols, 2023) at a semantic similarity threshold of 0.9. Pathway enrichment analysis was conducted using Enrichr, (Jawaid, 2025) querying the Reactome, KEGG, and WikiPathways databases, with pathways ranked by combined score. All statistical analyses and data visualization, including volcano plots, heatmaps, PCA, and enrichment bar plots, were performed in R (version 4.4.2) using R-package ggplot2 and pheatmap.

#### Kinase prediction analysis

Kinase activity prediction was performed using PhosR, (Kim *et al*., 2021a) integrating motif-based scoring and profile-based scoring. Protein sequences were retrieved via the UniProt API, and phosphosites with significant differences between normal and UL groups (paired t-test, *p* ≤ 0.05) were selected for further analysis. Kinase-substrate scoring was conducted using PhosphoSitePlus-based kinase-substrate relationships (Hornbeck *et al*, 2011) to identify key kinases involved in UL phosphosignaling.

### Immunohistochemistry

Normal myometrium tissue and UL tissue were sectioned at 3 μm on adhesive coated slides and stained with primary antibodies, PAGE4 (Abcam, ab224454), progesterone receptor (PR; Leica Biosystem, PGR-312-L-CE) and estrogen receptor (ER; Leica Biosystem, ER-6F11-L-CE). Briefly, after deparaffinizing in xylene and rehydration in graded alcohols (100%, 95%, 70%), endogenous peroxidase activity was blocked using 3% H2O2 (Fluka analytical, 95321-100ML) in methanol for 30 minutes and washed in two changes of phosphate buffered saline (PBS). Antigens were retrieved in microwave for 10 minutes in citrate buffer (pH6; Sigma-Aldrich, C9999) and then incubated for 1 hour at room temperature with 10% normal horse serum. Primary antibodies were diluted with antibody diluent (Agilent Technologies, S080983-2) and applied to the sections for overnight at 4°C with various dilutions (PAGE4, 1:500; ER, 1:10; PR, 1:100) in a humidifier dark chamber (Thermo Scientific, 44-0404-10). The sections were washed with three changes of PBS+Tween®20 and incubated with HRP-conjugated secondary antibodies (Fisher Scientific, NC9226446) for 30 minutes at room temperature. After rinsing in PBS+Tween twice, antigen-antibody complexes were visualized using Vector® NovaRed™ for 3 minutes. Finally, sections were counterstained with Meyer’s hematoxylin for 15 seconds and mounted using Krystalon.

#### Imaging and quantification

Slides were imaged using a Zeiss AxioImager microscope at the Light Microscopy Unit, University of Helsinki (LMU Bioimaging Services) using ZEN software (Zeiss) under standardized settings. Quantification of staining intensity was performed using ImageJ, where the staining area percentage (%) was calculated per sample.

#### Statistical analysis and visualization

All statistical analyses, data processing and visualization were conducted in R (version 4.4.2). Paired two-sample t-tests were performed to compare staining intensities between UL and normal groups for PAGE4, ER, and PR, with p-values calculated for each antibody.

### Affinity purification and proximity labeling

#### Site-directed mutagenesis and generation of MED12 and PAGE4 entry clones

The entry clone for wild-type PAGE4 and MED12 were obtained from the ORFeome collection (ORFeome and MGC Libraries; Genome Biology Unit, supported by HiLIFE, the Faculty of Medicine, University of Helsinki, and Biocenter Finland). Site-directed mutagenesis was performed to generate phospho-mimicking and phospho-deficient PAGE4 variants, as well as the *MED12* G44D variant, using primers listed in Table S1. Following amplification, the PCR product was treated with 0.5 µl of DpnI to digest the methylated template plasmid DNA, followed by overnight incubation at 37°C. The digested product was then transformed into *Escherichia coli*, and colonies were screened with kanamycin for PAGE4 and ampicillin for MED12. Plasmids were extracted using the NucleoSpin Plasmid Miniprep Kit (Macherey-Nagel) and validated through gel electrophoresis and DNA sequencing (Eurofins Genomics).

#### Cloning of PAGE4 and MED12 phosphovariants to MAC3-tag gateway destination vector

After the site-directed mutagenesis, the cDNA constructs were cloned into C terminal MAC3-tag (Liu *et al*, 2023; Liu *et al*, 2018; Liu *et al*, 2020) destination vector using Gateway™ LR Clonase™ Enzyme Mix (Life Technologies, 11791043). After *E.coli* transformation, expression clones were purified in the same manner as described previously. (Liu *et al*., 2023; Liu *et al*., 2018; Liu *et al*., 2020)

#### Generation of PAGE4 and MED12 Stable Inducible Cell Lines

All procedures were performed as previously described, with minor modifications. (Liu *et al*., 2020) In brief, Flp-In™ 293 T-Rex cells (Invitrogen, cat. no. R78007) were used to generate stable cell lines expressing MAC3-tagged constructs under a tetracycline-inducible system, ensuring single-copy transgene integration. Cells were cultured in low-glucose, tetracycline-free DMEM (Sigma-Aldrich) supplemented with 10% fetal bovine serum (FBS; Life Technologies, cat. no. 10270106) and 1% penicillin-streptomycin (Life Technologies, cat. no. 15140130) at 37°C in a humidified incubator with 5% CO₂. The cells were subsequently co-transfected with PAGE4 and MED12 expression vectors and the pOG44 Flp-Recombinase Expression vector (Invitrogen, cat. no. V600520) using Fugene6 transfection reagent (Promega, cat. no. E2691). Twenty-four hours post-transfection, cells were selected with 100 mg/ml hygromycin B (Life Technologies, cat. no. 10687010) for two weeks, after which positive clones were pooled and expanded, as described previously. (Liu *et al*., 2020) Stable cell lines were grown to 80% confluence, with nine 150 mm culture plates dedicated to AP-MS and another nine for BioID approach. Cells from three fully confluent plates were combined into a single biological sample, with a total of three biological replicates generated per stable cell line for each approach.

#### Sample processing

For AP-MS, cells were cultured until they reached 80% confluence, then induced with 2 μg/ml tetracycline for 24 hours before harvesting. For BioID, an additional 50 μM biotin was added alongside tetracycline 10 minutes prior to collection. Before harvest, samples were washed with wash buffer, collected using harvesting buffer, then pelleted by centrifugation, snap-frozen in liquid nitrogen, and stored at −80°C for further analysis. Green fluorescent protein (GFP) tagged with the MAC3-tag was used as a negative control and was processed in parallel with experimental samples. All the procedures were performed as described by Liu *et al*. (Liu *et al*., 2020)

#### Mass spectrometry

For various PAGE4 and MED12 stable cell lines, the desalted peptide samples were analyzed using the Evosep One liquid chromatography system coupled to a hybrid trapped ion mobility quadrupole time-of-flight (TIMS-TOF) mass spectrometer (Bruker timsTOF Pro/Pro 2) via a CaptiveSpray nano-electrospray ion source. Peptides were separated using an 8 cm × 150 μm column packed with 1.5 μm C18 beads (EV1109, Evosep) following the 60 samples per day (60-SPD) method, with a 21-minute gradient time. Mobile phases consisted of buffer A (0.1% formic acid in water) and buffer B (0.1% formic acid in acetonitrile).

Mass spectrometry analysis was conducted in positive-ion mode using data-dependent acquisition (DDA) in Parallel Accumulation Serial Fragmentation (PASEF) mode, with ten PASEF scans per TopN acquisition cycle. The DDA-PASEF-short_gradient_0.5s-cycletime method was applied for all AP-MS and BioID samples.

Raw data (.d files) were processed using FragPipe v21.1 with MSFragger-4.0, searching against the human protein fasta database from UniProtKB (downloaded on January 5, 2024), containing 40,954 entries (including 20,477 decoys, 50%). Carbamidomethylation of cysteine residues was set as a static modification, while N-terminal acetylation, methionine oxidation, and biotinylation of lysine and N-termini were considered variable modifications. Trypsin was selected as the cleavage enzyme, allowing for up to two missed cleavages. Peptide-to-spectrum match (PSM) values were reported for peptides with a false discovery rate (FDR) below 0.01, as determined by Philosopher. Label-free quantification parameters and instrument settings remained at their default values.

#### GO enrichment and statistical analysis

For protein-protein interaction (PPI) data obtained from stable Flp-In™ 293 T-Rex cell lines, intensity values were first imputed using quantile regression-based imputation for left-censored data (QRILC), as implemented in the imputeLCMD R-package. (Lazar & Burger, 2015) Median normalization was applied across samples to adjust for variations in protein abundance, followed by log2 transformation to improve data distribution.

High-confidence interactors (HCIs) were identified using Significance Analysis of INTeractome (SAINT) express version 3.6.0 and Contaminant Repository for Affinity Purification (CRAPome, http://www.crapome.org/) were used for additional filtering. Proteins achieving a SAINT score ≥ 0.85 were retained, corresponding to an estimated Bayesian false discovery rate (BFDR) of ≤ 0.05. Further filtering was applied using CRAPome, excluding proteins with a contaminant frequency ≥ 20%, while only interactors with an average spectral count fold change ≥ 3 were classified as HCIs.

Gene Ontology (GO) enrichment analysis was performed using clusterProfiler, separately analyzing Biological Process (BP), Cellular Component (CC), and Molecular Function (MF). To reduce redundancy, enriched GO terms were refined using rrvgo with a similarity threshold of 0.9. Pathway enrichment analysis was conducted using Enrichr, querying Reactome, KEGG, and WikiPathways databases, with pathways ranked by combined score. All data visualization was performed using R (version 4.4.2).

### Luciferase reporter assay

For luciferase pathway analysis, HEK293 cells were seeded in 96-well plates at a density of 2 × 10⁵ cells/ml and allowed to adhere. The following in day, cells were then transfected with 50 ng of PAGE4 and MED12 plasmids, 47.5 ng of Firefly luciferase reporter plasmid containing the promoter of interest, and 2.5 ng of Renilla luciferase plasmid as an internal control. Transfections were carried out using Fugene6 transfection reagent following the manufacturer’s instructions, and cells were incubated at 37°C with 5% CO₂ for 24 hours. Once cells reached 80-90% confluency, they were processed using Dual-Luciferase® Reporter Assay System (Promega, E1910). Luciferase activity was measured on a CLARIOstar luminometer (BMG LABTECH, Ortenberg, Germany) at a gain setting of 3900. Firefly luciferase signals were normalized to Renilla luciferase to account for transfection efficiency, and data were analyzed as relative luciferase units (RLU). Data were analyzed and plotted using GraphPad Prism (version 9.4.1, GraphPad Software, San Diego, CA, USA). Statistical significance was determined using one-way ANOVA, with a significance threshold set at *p* < 0.05. Data are presented as mean ± standard deviation (SD).

### Chromatin immunoprecipitation sequencing (ChIP-seq)

ChIP-Seq assays were performed as previously described, with modifications to optimize experimental conditions. (Gawriyski *et al*, 2024) In brief, HEK293 cells were seeded in 10 cm culture dishes, and after 24 hours, tetracycline was added to induce the expression of PAGE4 and MED12 proteins with the final concentration of 2 μg/ml. Cells were cross-linked with 1% formaldehyde for 10 minutes at room temperature, followed by quenching with 125 mM glycine. After two washes with pre-chilled PBS, cells were lysed in hypotonic lysis buffer (20 mM Tris-Cl, pH 8.0, 10% glycerol, 10 mM KCl, 2 mM DTT, and a protease inhibitor cocktail (Roche)) to isolate nuclei. The nuclear pellets were washed in cold PBS and lysed in a 1:1 mixture of SDS lysis buffer (50 mM Tris-HCl, pH 8.1, 1% SDS, 10 mM EDTA, and protease inhibitors) and ChIP dilution buffer (16.7 mM Tris-HCl, pH 8.1, 0.01% SDS, 1.1% Triton X-100, 1.2 mM EDTA, 167 mM NaCl, and protease inhibitors). Chromatin was sheared to an average fragment size of ∼300 bp using a Q800R sonicator (QSonica) at 4 °C. Dynabead Protein G (Invitrogen) was pre-washed in blocking buffer (0.5% BSA in IP buffer: 20 mM Tris-HCl, pH 8.0, 2 mM EDTA, 150 mM NaCl, 1% Triton X-100, and protease inhibitors) and incubated with anti-POLR2 antibody (RPB1 CTD, clone 4H8, Cell Signaling Technology, #2629). The prepared chromatin lysate was incubated with antibody-coated beads for 12 hours, followed by four washes with washing buffer (50 mM HEPES, pH 7.6, 1 mM EDTA, 0.7% sodium deoxycholate, 1% NP-40, 0.5 M LiCl) and two washes with 100 mM ammonium hydrogen carbonate (AMBIC) solution. DNA-protein complexes were eluted in extraction buffer (10 mM Tris-HCl, pH 8.0, 1 mM EDTA, 1% SDS), and proteinase K with NaCl was added to reverse the cross-links. DNA was purified using a Mini-Elute PCR Purification Kit (Qiagen). The NEBNext® Ultra™ II DNA Library Prep Kit (E7103L, NEB) was used for ChIP-Seq library preparation, and the resulting libraries were sequenced on an Illumina NovaSeq 6000 S4 sequencing system (Illumina, San Diego, CA, USA).

### ChIP-Seq library processing and analysis

ChIP-seq libraries were sequenced to yield 150 bp paired-end reads. Quality control was performed with FastQC (v0.11.9) to assess read integrity. Subsequently, TrimGalore (v0.6.7, RRID: SCR_011847) was used to remove adaptor sequences and discard reads below the length cutoff. The resulting high-quality reads were aligned to the hg38 human reference genome via Bowtie2 (v2.2.5) (Langmead & Salzberg, 2012) under default parameters. SAMtools (v1.9) (Danecek *et al*, 2021) was then employed to eliminate low-quality alignments using “-q 30 -F 3844.” Duplicate reads were identified and removed with the Picard toolkit (v2.25.1, RRID: SCR_006525). Peak detection was carried out using MACS2 (v2.1.4) (Zhang *et al*, 2008) at a p-value threshold of <1×10^⁻³^, and the HOMER script “annotatePeaks.pl” (v4.11.1) (Heinz *et al*, 2010) was employed to annotate the resulting peaks.

### Data and code availability

All R scripts, processed data, and raw source tables used in this study are publicly available on GitHub at https://github.com/bongrita/MED12_PAGE4_multiomics. The mass spectrometry dataset generated in this study is available from the MassIVE database (https://massive.ucsd.edu/) with web access MSV000101051(Password: Varjosalo). This repository is archived on Zenodo for long-term access and citation at: https://doi.org/10.5281/zenodo.15574453.

## Supplementary Figures

**Figure S1.** Protein and RNA expression profiles of PAGE4 in human reproductive organs. based on sex Tissue-specific RNA and protein expression levels (normalized transcripts per million, nTPM) in male and female reproductive organs. Expression data were obtained from the Human Protein Atlas (https://www.proteinatlas.org/). The upper panel shows female tissues, and the lower panel shows male tissues. Bars indicate RNA (light colors) and protein (dark colors) expression levels, highlighting sex-specific tissue expression patterns.

**Figure S2.** Functional enrichment of differentially expressed proteins across UL genotypes. GO enrichment analysis of significantly up- and down-regulated proteins identified from DIA-based proteome profiling of UL and matched MM samples. Differential expressions were analyzed separately for each genetic subtype, (A) MED12, (B) COL4A5-COL4A6, (C) HMGA2, and (D) FH with significantly changing proteins (p < 0.05) divided into up- and down-regulated sets. For each group, the left panels display the top enriched biological process cellular compartments, and molecular function GO terms (bubble size = number of proteins; color = adjusted p-value), while the right panels show top 10 corresponding pathway categories plotted by enrichment significance (-log10 p-value).

**Figure S3.** Pathway enrichment analysis of PAGE4 phosphomutants and MED12 interactomes. Heatmap depicting significantly enriched pathways identified through enrichment analysis across five selected PAGE4 and MED12 baits, highlighting key biological processes including transcription regulation, DNA repair, RNA processing, cell cycle control, and cancer-related signaling pathways. Enrichment terms were filtered based on combined score ranking and occurrence across multiple baits to identify shared and distinct interactome functions.

**Figure S4.** Luciferase reporter assays for pathway activation across PAGE4 variants and MED12 mutants. Relative luciferase activity (Firefly/Renilla) for PAGE4 WT, PAGE4 phosphomutants, MED12 WT, and MED12 G44D across six key signaling pathways: (A) cell cycle/pRb-E2F, (B) p53/DNA damage response, (C) MAPK/ERK, (D) Wnt signaling, (E) estrogen receptor, and (F) MAPK/JNK. Data represent mean ± SEM from at least three independent experiments. Statistical significance was determined using one-way ANOVA with multiple comparisons.

**Figure S5.** ChIP-seq analysis of POLRII binding across PAGE4 and MED12 constructs. (A) Peak intensity distributions reflect signal strength variability across samples. (B) Peak genomic distribution highlights consistent promoter enrichment, with subtle shifts in intronic and intergenic engagement in PAGE4 phosphomutants.

